# Morph-specific plasticity drives divergent host-shift and acclimation strategies in aphids

**DOI:** 10.64898/2026.02.11.705319

**Authors:** Rituparna Ghosh, Aurélie Etier, Maxime Berberon, Nathalie Marnet, Solenne Berardocco, Alain Bouchereau, Jean-Christophe Simon

## Abstract

Understanding how organisms cope with environmental change is central to eco-evolutionary theory, yet a key unresolved challenge is distinguishing anticipatory plasticity, where phenotypes are pre-configured using predictive cues, from responsive plasticity, where phenotypes are adjusted after environmental exposure. Empirically testing these strategies is challenging because phenotypes measured after environmental transitions integrate both prior cue-dependent state configuration and post-exposure adjustment, obscuring when plastic responses are initiated. Systems with predictable environmental transitions and discrete life-history stages provide a powerful opportunity to resolve this challenge. Here, we use host-alternating aphids to test how plasticity strategies are organized across morphs experiencing the same seasonal host shift but facing different ecological constraints. We quantified fitness, transcriptomic responses, and host chemical landscapes across three morphs transferred among secondary hosts differing in phloem composition. Migratory spring morphs maintained uniformly high survival and showed minimal transcriptional change, consistent with anticipatory plasticity mediated by pre-configured metabolic and detoxification programs. In contrast, summer morphs exhibited host-dependent survival costs and extensive regulatory remodeling, indicating responsive plasticity. By integrating fitness, transcriptomics, and metabolomics, we show that predictable environmental transitions can favor the evolution and partitioning of plasticity strategies across life-history stages. More broadly, our results provide mechanistic evidence linking environmental predictability, regulatory architecture, and the evolution of plastic responses to heterogeneous environments.

## Introduction

Natural environments are heterogeneous across space and time, exposing organisms to predictable seasonal changes and unpredictable variation in resources, abiotic conditions and biotic interactions ^1,2^. A widespread evolutionary response is the evolution of complex life cycles, in which a single genotype produces different life stages that exploit contrasting ecological niches over time ^3–5^. Such life cycles occur across animal taxa, including parasites cycling among hosts ^6^, amphibians transitioning between aquatic and terrestrial habitats ^7^, and insects alternating seasonally between habitats, resources or dispersal modes ^8^. In these systems, phenotypic plasticity is essential: individuals must match physiology, behaviour and often morphology to environments that differ sharply, sometimes over short timescales, and selection acts not only on trait values but on the mechanisms enabling reliable transitions ^9^.

Phenotypic plasticity is itself an evolvable trait subject to selection ^4,10^. Its magnitude, timing, and regulatory mechanisms can vary among genotypes, populations and life-history stages, and plastic responses can be adaptive, neutral, or maladaptive depending on environmental predictability and the costs of phenotype switching ^11–14^. A key conceptual distinction can be drawn between *anticipatory plasticity*, in which phenotypic adjustments occur before an environmental change based on reliable cues, and *responsive plasticity*, in which phenotypes are modified after exposure to new conditions ^12,15–17^. Anticipatory plasticity can reduce the immediate fitness costs of abrupt environmental transitions by pre-configuring physiological or regulatory states in advance, but it requires cue reliability and may impose maintenance costs if pre-activated pathways are unnecessary ^18^. Responsive plasticity may be more flexible, because it tracks the current environment directly, but it can entail delays during which individuals experience stress, mortality or reduced reproduction while regulatory programs adjust ^19,20^. Despite their fundamental importance in eco-evolutionary theory, the mechanistic basis of anticipatory versus responsive plasticity in natural systems remains poorly resolved. This is largely because most empirical studies measure plasticity only after environmental change, obscuring whether observed phenotypes reflect pre-configured states or active post-exposure adjustment ^12,17^. Distinguishing these strategies requires systems in which distinct life-history stages or morphs experience the same predictable environmental transitions but differ in their ecological roles and in the selective pressures shaping their capacity for acclimation. Such systems allow plasticity strategies to be inferred by contrasting phenotypes expected to rely differentially on anticipatory versus responsive regulation.

Herbivorous insects provide particularly tractable systems to address this question because their performance is tightly constrained by host plant chemistry, and host use often varies predictably across life stages ^21^. Aphids (Hemiptera: Aphididae) are emblematic in this respect. They exhibit cyclical parthenogenesis, alternating between asexual and sexual reproduction, and striking polyphenisms in which a single genotype can give rise to discrete alternative phenotypes in response to environmental cues ^22^. Because they can be propagated as clonal lines under laboratory conditions, aphids constitute powerful systems for studying phenotypic plasticity, as their genetic uniformity reduces confounding genetic effects and enables a direct assessment of environmentally induced phenotypic variation. Aphids have therefore emerged as model organisms for investigating the genetic and epigenetic bases of phenotypic plasticity, including wing dimorphism, variation in reproductive mode, and host-plant acclimation ^8,22–28^. As a striking example of this phenotypic plasticity, approximately 10% of aphid species exhibit seasonal host alternation between a woody primary host, used for overwintering and sexual reproduction, and one or more herbaceous secondary hosts exploited during the growing season ^29,30^. These hosts are often taxonomically unrelated and differ sharply in nutrient composition and defence chemistry, implying that host alternation relies on rapid acclimation. Acclimation to new hosts in aphids is known to involve coordinated chemosensation, regulation of feeding behaviour, salivary secretions, nutrient transport and assimilation, and detoxification of plant metabolites, and these responses can be mediated by transcriptional, post-transcriptional and potentially chromatin-based regulation ^25,31–37^.

Selective pressures associated with host acclimation are unlikely to be uniform across the aphid life cycle, particularly in host-alternating (heteroecious) species where seasonal migration links two ecologically and taxonomically distinct hosts ^31,38–40^. In heteroecious species, spring migrants (SM) that initiate colonization of secondary hosts face the most stringent challenge: they must migrate, rapidly settle and establish feeding on unfamiliar plants following an effectively irreversible host shift, as they are no longer able to feed on the primary host they leave to reach the secondary host; failure results in immediate mortality or severe reproductive loss ^39,41,42^. In contrast, summer morphs develop on already accepted secondary hosts and typically experience reduced exposure to novelty. Wingless summer morphs (WlSM) are sedentary and specialized for local population growth, whereas winged summer morphs (WgSM) retain dispersal capacity and may occasionally encounter novel hosts within the secondary host range, suggesting intermediate selection for broad tolerance and fast acclimation ^43^. Such life-stage–specific contrasts may favour distinct plasticity strategies among morphs: migratory morphs are expected to rely on anticipatory plasticity, expressing pre-configured physiological and regulatory states that buffer survival at host transfer, whereas resident morphs should depend more strongly on responsive plasticity, adjusting gene expression after settlement and potentially with greater costs and host-dependent failures.

Here we test this hypothesis using the bird cherry–oat aphid, *Rhopalosiphum padi*, a globally important cereal pest that alternates between a single woody primary host (*Prunus padus,* Rosaceae) and a broad range of grass (Poaceae) hosts ^41,44,45^ (Extended data fig. 1a). We compared the plastic responses of three morphs (SM, WlSM and WgSM) after transfer to four Poaceae hosts differing in nutritional and chemical composition, by quantifying survival, fecundity, and whole-transcriptome responses in each morph (Extended data fig. 1b). To link host quality to the mechanisms underlying morphs’ plastic responses to plant shifts, we combine transcriptomics with targeted phloem metabolomics and experimentally manipulate a metabolite that strongly differentiates hosts. This integrated design allows us to test whether predictable host transitions favour morph-specific plasticity strategies, including anticipatory versus responsive plasticity, and to identify the regulatory architectures and chemical drivers associated with acclimation strategies.

## Materials and methods

### Insect and plant sources

In early March 2023, ten *Rhopalosiphum padi* fundatrices (first morph of the annual life-cycle, hatching from winter eggs) of the age of first two larval developmental stages were collected from *Prunus padus* trees in Le Rheu, France (48.1011° N, 1.7954° W). Each individual was placed in individual clip cages (1.5 cm in diameter) on young leaves to prevent overcrowding, which could impact their ability to detect plant cues. The cages were attached to the underside of uninfested leaves. Spring wingless morphs produced by fundatrices were reared on mature *P. padus* leaves (30-days old plants as determined in Etier *et al*., 2025^42^ to induce spring migrant (SM) production. These morphs were either installed directly on different secondary hosts—spring oat (*Avena sativa*, ‘Céleste’ cultivar), barley (*Hordeum vulgare*, ‘KWS Faro’ cultivar), Italian ryegrass (*Lolium perenne*, ‘Syntilla’ cultivar) and wheat (*Triticum aestivum, “Geny”* cultivar)—for survival tests and transcriptomics. Some SM individuals were also installed on a fifth secondary host, orchard grass (*Dactylis glomerata*), to produce wingless summer morph (WlSM) and winged summer morph (WgSM) on a common host prior to host shifting. This fifth host was used to avoid conditioning effects. WlSM and WgSM were then transferred from orchard grass to one of the four previously listed hosts and used for survival assays and transcriptomic analyses.

*P. padus* saplings, measuring 75–100 cm in height, were sourced from local nurseries and planted separately in 3-liter pots filled with soil (60% white peat, 40% sand, 14% coconut fibre medium, 6% clay). Additionally, seeds of five grasses were individually germinated in 15 mL plastic tubes with drainage holes. 15-day-old seedlings were used in experiments. No fertilizers or pesticides were applied to plants. These plants were used for aphid rearing, aphid fitness assays, and metabolomics. Both the plants and aphids were maintained in a controlled climatic chamber with constant growth conditions (temperature- 18°C, photoperiod- 16-hour light, 8-hour dark cycle).

### Aphid survival tests

Survival of SM, WlSM, and WgSM was assessed on the four secondary host species. For SM, individuals that were 48 hours old post-adult emergence (after the final molt) and actively flying (collected from the cage containing *P. padus* saplings) were used for survival experiments. Each aphid was placed individually on a secondary host plant and enclosed within a perforated plastic bag to allow gaseous exchange. A total of 20 replicates per host species were conducted. WlSM and WgSM were collected from orchard grasses and subjected to survival tests under the same conditions as the SM. Survival was recorded every alternate day for nine consecutive days. In addition to survival tracking, the reproductive output of each morph was recorded over the nine days by counting the number of offspring produced per individual, providing further insights into aphid fitness.

### Aphid RNA extraction and sequencing

Adults of the SM morph were sampled 72 h after reaching adulthood, following 24 h on the primary host *P. padus* and a subsequent 48 h on one of four secondary hosts (oat, barley, wheat, or ryegrass). Adults of the WlSM and WgSM morphs were sampled 72 h after reaching adulthood, following 24 h on *D. glomerata* and a subsequent 48 h on one of the same four secondary hosts. Eight individuals were pooled per biological replicate, with four replicates per condition, except for WlSM on ryegrass (three replicates due to high mortality). Samples were immediately flash frozen in liquid nitrogen prior to RNA extraction.

Total RNA was extracted using the RNeasy Mini Kit (Qiagen, cat. no. 74104) with an additional grinding and 2 min incubation at 56 °C before homogenization on Qiashredder columns (Qiagen, cat. no. 79656). RNA quality was assessed by NanoDrop 2000 and Agilent 2100 Bioanalyzer (RIN > 6). Strand-specific paired-end poly(A)-selected libraries (n = 47) were prepared by Genewiz (Azenta Life Sciences) and sequenced on an Illumina NovaSeq platform (2 × 150 bp), generating ∼40 million paired-end reads per sample (≥ 80 % bases Q ≥ 30).

Reads were processed using the nf-core/rnaseq pipeline v3.12 (Nextflow 22.10.4; https://nf-co.re/rnaseq/3.12.0), with adapter trimming (TrimGalore v0.6.7 / Cutadapt v3.4) and quality control (FastQC v0.11.9, MultiQC v1.13). Reads were aligned to the *Rhopalosiphum padi* reference genome (assembly ASM2088224v1; RefSeq accession GCF_020882245.1) using STAR v2.7.9a. In addition to the standard RefSeq gene models, the annotation was augmented with functionally relevant gene sets, including candidate salivary effector genes inferred from orthology with *Acyrthosiphon pisum*. These effectors were classified into three categories: (i) “expressed effectors”, expressed in salivary glands; (ii) “up-effectors”, upregulated in salivary glands relative to gut tissue; and (iii) “salivary proteins”, identified by proteomic analyses ^46^. In addition, odorant and gustatory receptor annotations specific to *R. padi* ^47^, as well as bicycle genes identified using a published classifier ^48^, were also included. Alignment outputs were used to assess quality control metrics including mapping rate, duplication, and rRNA content. Transcript-level quantification was performed using Salmon v1.10.1 in quasi-mapping mode, and gene-level counts were summarized with tximport/tximeta (Bioconductor v1.12.0 / 1.24.0).

Raw read counts were imported into the AskoR framework for differential expression analysis. Counts were normalized using the TMM (Trimmed Mean of M-values) method, and differential expression was assessed with the edgeR quasi-likelihood pipeline. Genes with an adjusted *P*-value < 0.05 (Benjamini–Hochberg correction ^49^) and |log₂ fold change| ≥ 1 were considered significantly differentially expressed. To complement the model-based approach and account for potential deviations from negative-binomial assumptions, a Wilcoxon rank-sum test was performed in parallel on normalized expression values with the same thresholds as for EdgeR: *P*-value < 0.05 and |log₂ fold change ≥ 1. This non-parametric procedure allowed the detection of expression shifts that may not be captured by distribution-based statistical models, as already shown by ^50^. Genes identified as significant by either method were retained, and the final set of differentially expressed genes corresponded to the union of edgeR-derived and Wilcoxon-derived gene lists.

Clustering analyses were conducted on the 1,500 genes with the highest median absolute deviation (MAD) across samples. Co-expression clusters were identified using a consensus clustering approach to obtain robust gene groupings across resampling iterations. Gene Ontology enrichment was then assessed for each cluster using the topGO package (weight01 algorithm). GO terms with Benjamini–Hochberg–corrected *P*-values < 0.05 were considered significantly enriched. The background genes consisted of all expressed genes retained after low-expression filtering (≥ 3 counts per million in at least half of the samples; 9,811 genes). Enrichment tests compared the gene composition of each cluster against this filtered background set.

PCA and hierarchical clustering were performed to assess sample similarity and expression patterns.

### Phloem sample preparation

Phloem samples (n = 10) were collected from leaves of two-week-old plants of each tested secondary host (wheat, oat, barley and ryegrass) following the centrifugation method ^51,52^. 30 µl of phloem was mixed with 30 µl cold methanol containing 200 mM beta-aminobutyric acid (BABA) and 400 mM adonitol. The mixture was sonicated at 50 Hz for 3 minutes and centrifuged for 10 minutes at 3000 rpm at room temperature to remove proteins and particles. This mixture was divided into two parts for the amino acid and sugar work. All the samples were completely dried using a Speed-Vac concentrator at 35°C.

### UPLC-UV(DAD)-based analysis of amino acids of phloem samples

To the dried sample, 15 µl (same volume of solvent that was evaporated) of MilliQ water was added. For derivatization of amino acids, The AccQTag derivatization kit (Waters, Milford, MA, USA) was used according to the manufacturer’s protocol ^53,54^. For every batch of samples analyzed, a commercial mixture of amino acid standards (each amino acid with 100 µM concentration) was analyzed randomly between the samples to avoid instrument response-related variations. Amino acids were analyzed using Empower software (Waters). Peak area of amino acids was normalized by the peak area of the internal standard BABA. Absolute amino acid concentrations were calculated by comparing the normalized peak area of one amino acid to the peak area of that amino acid in the external standard where its concentration was known to be 100 µM.

### GC-MS(QQQ)-based analysis of sugars and sugar alcohols of phloem samples

To the dried sample, 35 µL (same volume of solvent that was evaporated) of pyridine containing methoxyamine chloride with a concentration of 20 g L^-1^ was added. To dissolve the pellet, the samples were vortexed for 10 seconds and incubated on a dry bath at 40°C for an hour. To that, 35 µl of the derivatizing reagent (trimethylsilyl)-trifluoroacetamide (MSTFA) was added and the mixture was incubated for another half an hour at 40°C. Immediately after derivatization process, 1 µL of sample was injected into a GC (GC-2010 Plus, Shimadzu) coupled with a triple quadrupole MS (GCMS-TQ8040, Shimadzu) with a split ratio 1:40 at inlet temperature 260°C. Metabolites were separated by a TG-5MS column (26098-1420, Thermo Scientific) (length- 30 m, inner diameter- 0.25 mm, and film thickness- 0.25 µm). Helium gas with a flow rate of 1 mL min^-1^ was used as carrier gas. Oven temperature program was followed as described in ^55^.

In the MS, the ionization was achieved by electron impact at 70 eV. A solvent delay of 5.65 min was maintained before MS acquisition was started. A Q3 scan mode with event time 0.2 sec and scan speed 5000 was maintained over the m/z range 45-950. Metabolite peaks were identified and processed using “GCMS Solutions” software (Shimadzu). Sugar identification was made by comparing the mass spectrum and retention time with an external standard mixture. For quantification, a standard curve method-based quantification was followed. Standard curves were prepared by injecting five external sugar standard-mixes, with each sugar having different concentrations (7.81, 15.63, 250, 500, and 750 µM).

### Organic acid complementation assay

To assess the role of organic acids in aphid performance, a compound complementation assay was performed using wheat. Quinic acid, highly abundant in the least preferred host (ryegrass)—was selected for the assay. Two-week-old wheat seedlings were excised at the stem–root junction, and the root portion was discarded to facilitate compound uptake via the cut stem.

Quinic acid solutions (2.5 mM) were prepared by dissolving the compounds in tap water. Individual cut seedlings were placed in 10 mL sterile Corning® culture tubes containing 3 mL of the solution. For each treatment (quinic acid, and water controls), thirty biological replicates were maintained. To ensure the consistent availability of the complemented compounds and to prevent degradation of the plant material, the test plants were replaced with freshly prepared seedlings twice a week. Following a 30-minute incubation of the plant in the respective solution, a single WlSM was introduced onto each seedling. The entire setup was enclosed within a transparent, perforated plastic bag to maintain humidity and allow aeration.

To control for any pH-induced effects resulting from the acidic nature of the solutions (pH 4.5–5.0), an additional acidic water control (tap water adjusted to the pH of the acid solutions) was compared with the standard water control. Aphid survival and fecundity were monitored at regular intervals throughout the experiment.

### Data analysis and statistics

Survival was analysed using a Cox proportional hazards model to assess the association between morph type (SM, WlSM, and WgSM) and time to death after transfer to the four test plants (barley, oat, wheat, and ryegrass). Survival analyses were performed using the *survival* package in R. Kaplan–Meier survival curves were generated to visualize survival dynamics over the assay period and to estimate survival probabilities for each morph–host plant combination.

Fecundity analyses were performed using only individuals that survived until the day of observation. Cumulative fecundity (number of offspring produced per living individual at day 2, 4, 7 and 9 after transfer) was modelled using a linear mixed-effects model with morph, host plant, and their interaction as fixed effects, and sample as a random intercept to account for repeated measures within individuals. The model was fitted with the *lmer* function of the lme4 package, and *P*-values for fixed effects were obtained using the lmerTest package. Post hoc comparisons were conducted using estimated marginal means (*emmeans*), with Tukey adjustment for multiple testing. Pairwise contrasts were computed for (i) morph, (ii) host plant, (iii) morph levels within each host plant, and (iv) host plant levels within each morph. To visualize cumulative fecundity dynamics over time, LOESS-smoothed curves were generated with *ggplot2* for each morph–host combination.

Data for metabolite concentrations (mean ± SE) were first assessed for variance homogeneity using Levene’s test. Variables meeting the assumptions of normality and homogeneity were analyzed using one-way ANOVA, followed by Tukey’s HSD post hoc test to identify pairwise differences (p ≤ 0.05). For datasets that violated parametric assumptions, Kruskal–Wallis tests were applied, and significant differences were evaluated using Dunn’s post hoc test with Bonferroni correction (p ≤ 0.05). Heatmap visualization, principal component analysis (PCA), and Random Forest classification were conducted using MetaboAnalyst 6.0 (https://www.metaboanalyst.ca/). Random Forest models were trained with 500 trees under default tuning parameters. Model robustness was evaluated using the out-of-bag (OOB) error rate, which provides an internal cross-validation estimate of classification accuracy. Variable importance was quantified using permutation-based mean decrease accuracy, which measures the decline in predictive performance when individual metabolites are randomly permuted while keeping all others unchanged. Ternary nutrient plots were constructed in PAST v4.03 ^56^ using normalized proportions of pooled amino acids, sugars, and organic acids.

### Data availability

Raw RNA-seq data and processed count matrices will be deposited in the NCBI GEO or ENA database under accession number [to be provided upon acceptance].

## Results

### Morph-specific survival and fecundity across secondary hosts

To test whether morphs with contrasting dispersal ecology (SM, WlSM, WgSM) exhibit distinct acclimation outcomes under novel host environments, we quantified survival and reproductive output following transfer to four Poaceae hosts (wheat, barley, oat, and ryegrass). Survival was monitored for nine days after transfer from *P. padus* (SM), or from a common natal host (orchard grass, *Dactylis glomerata*; summer morphs) to each secondary host, capturing immediate fitness consequences of host switching (Fig. 1a–f). SM exhibited uniformly high survival on all hosts, with no host-specific differences in mortality risk (Fig. 1a, Supplementary table 1), consistent with broad tolerance during the colonization phase. In contrast, both summer morphs showed host-dependent survival. WlSM experienced elevated mortality on ryegrass relative to oat (HR = 1.80, *p* = 0.03) and wheat (HR = 1.93, *p* = 0.02) (Fig. 1b). WgSM showed an even more pronounced pattern. Ryegrass imposed a strong survival cost relative to barley (HR = 2.67, *p* = 0.006), whereas survival on wheat and oat was intermediate and did not differ significantly from other conditions (Fig. 1c). The survival penalties in summer morphs emerged early after transfer, consistent with acute establishment stress on poorer hosts. Fecundity further differed among morphs. SM displayed pronounced host-dependent variation in reproductive output: highest on wheat and oat and reduced on barley and ryegrass (oat vs ryegrass: estimate = 4.58, *p* < 0.001; wheat vs ryegrass: estimate = 4.68, *p* < 0.001; oat vs barley: estimate = 5.35, *p* < 0.001; wheat vs barley: estimate = 5.44, *p* < 0.001) (Fig. 1d; Supplementary table 2). In contrast, WlSM showed no significant differences in fecundity across hosts among survivors (all *p* > 0.7; Fig. 1e, Supplementary table 2), implying that poor hosts primarily reduce fitness via survival rather than reproduction once feeding is established. WgSM exhibited no differences in fecundity among barley, wheat, and oat (all *p* > 0.11), although a non-significant trend toward reduced reproduction on oat was observed (Fig. 1f), Supplementary table 2). A complete reproductive failure on ryegrass indicates a more severe colonization constraint in this morph when exposed to the least suitable host. Together, SM combine host-general survival with host-dependent reproductive investment, whereas summer morphs exhibit host-specific survival constraints and, in WgSM, a reproductive collapse on ryegrass.

**Fig. 1:**
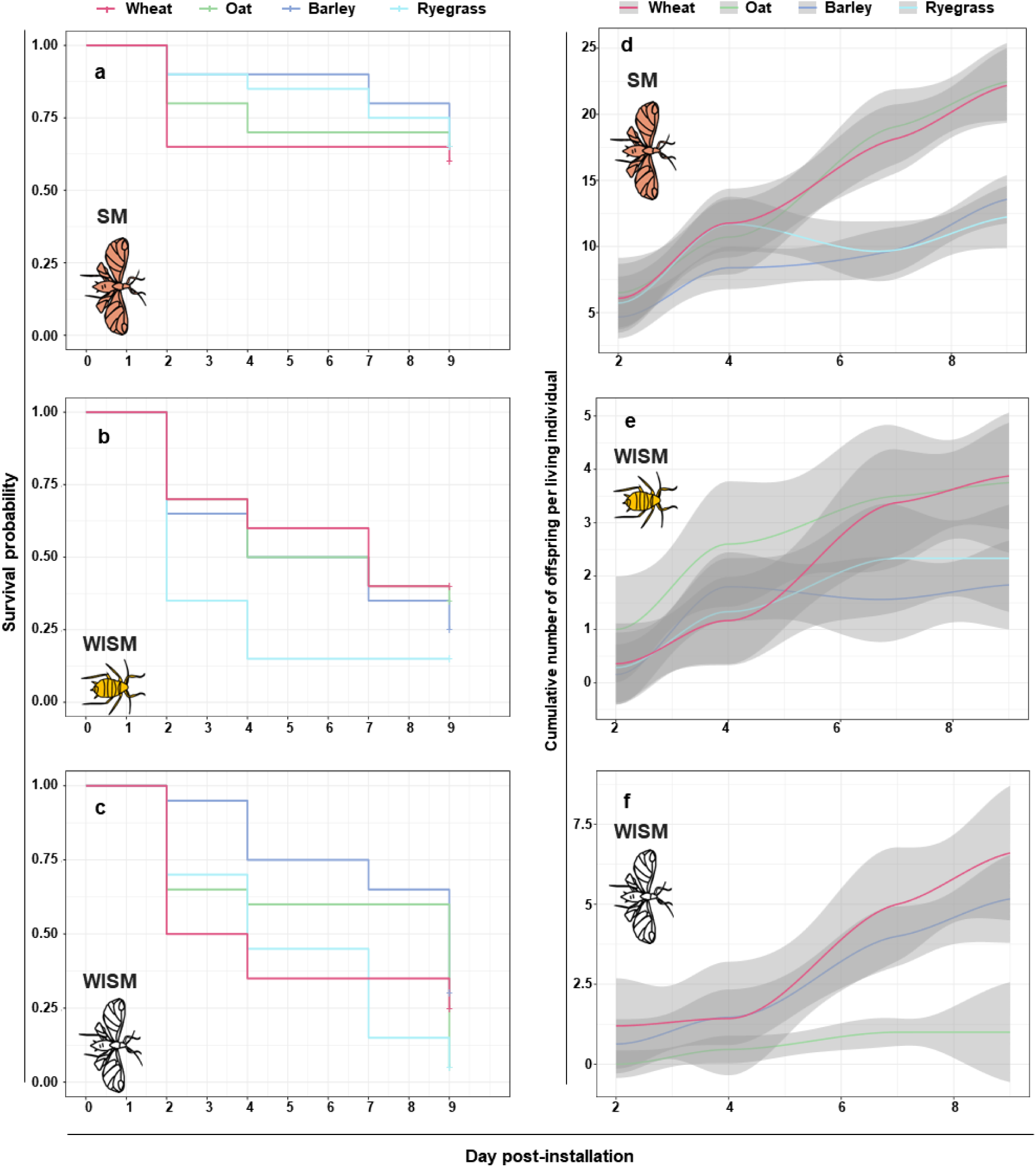
Survival and fecundity dynamics of three morphs (spring migrant, SM; wingless summer morph, WlSM; winged summer morph, WgSM) of the aphid *Rhopalosiphum padi* on four host plants (barley, oat, ryegrass, and wheat). Panels (a–c) show Kaplan–Meier survival curves, and panels (d–f) show cumulative fecundity curves per surviving adult. Each morph was analyzed independently. Shaded areas indicate 95% confidence intervals, and LOESS smoothing was applied to fecundity data for visual clarity.

### Transcriptional responses to host switching are structured by morph identity

To determine whether fitness differences reflect distinct gene regulatory responses, we performed whole-transcriptome analyses of each morph following host transfer. Across analyses, morph identity dominated variation (Fig 2a-c). Principal component analyses separated SM from summer morphs along the primary axis (34.9% variance) and distinguished WlSM and WgSM along the secondary axis (13.2%). Within morphs, host-associated clustering was weak (Extended data fig. 2), consistent with gene expression profiles that were highly correlated across hosts within each morph, indicating largely morph-specific, and host-general global transcriptional programs. Differential expression analyses reinforced this pattern and revealed a gradient in responsiveness across secondary host comparison (Fig 2b-c). SM exhibited remarkably few differentially expressed genes among host-pairs, indicating minimal transcriptional divergence across secondary hosts (Supplementary table 3). WlSM showed modest non-significant host-responsive change, whereas WgSM exhibited a significant extensive differential expression across host contrasts, consistent with large-scale host-dependent remodeling after settlement (Supplementary table 3). Responsive genes were largely morph-specific, supporting the view that morph identity constrains both baseline expression states and the set of pathways available for host-associated modulation.

**Fig. 2:**
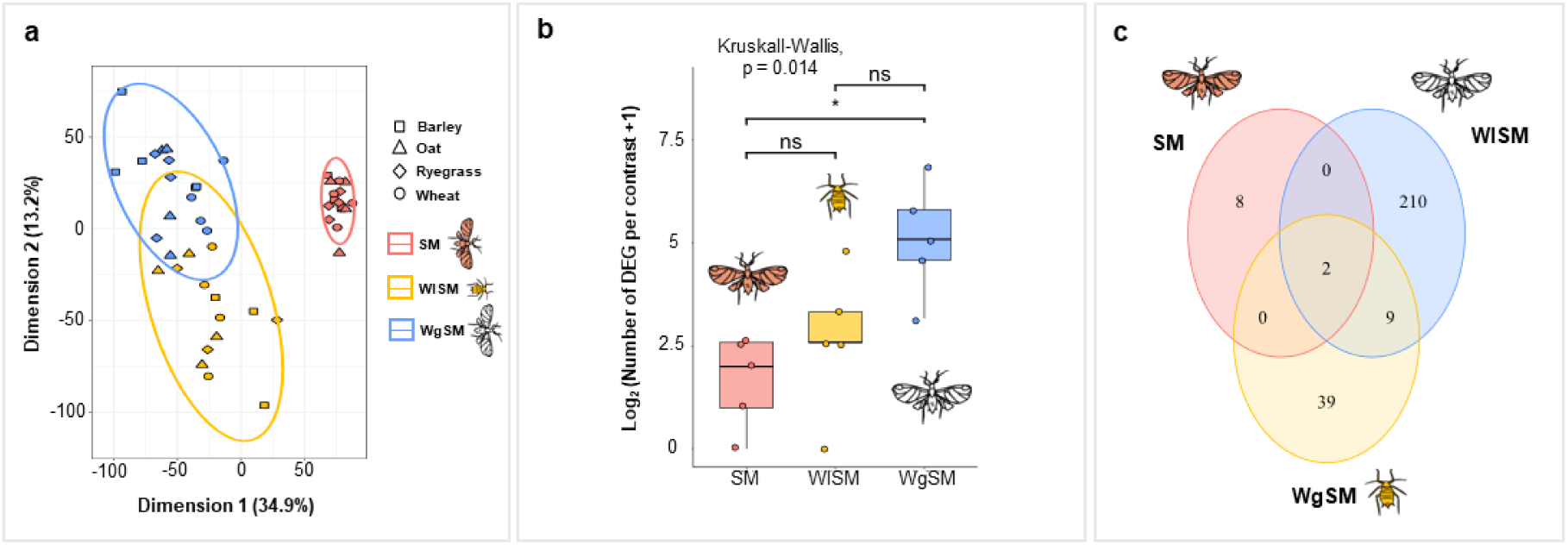
Morph-specific patterns of differential gene expression of *R. padi* in response to host shifts. **a,** Principal component analysis of all samples showing the first two principal components (PC1–PC2), with points colored by morph (SM, WlSM, WgSM). SM samples are clearly separated from summer morphs along PC1, whereas WlSM and WgSM are further distinguished along PC2. **b,** Boxplots showing the log2 transformed number of differentially expressed genes (DEGs) detected for each morph (SM, WlSM, WgSM) from the pairwise comparisons between all hosts. Each box represents the median and interquartile range (25ᵗʰ–75ᵗʰ percentiles) and points show individual values. A Kruskal-Wallis test was used to assess overall differences among morphs, followed by pairwise Wilcoxon tests with Bonferroni correction. Significant differences between morphs are indicated by asterisks (p < 0.05). **c,** Venn diagram shows the overlap of DEGs among SM, WlSM, WgSM after removing duplicate genes. Each circle represents the total number of DEGs for a given morph, and intersections indicate genes shared between morphs. Genes unique to a morph are shown outside the overlapping regions.

Functional annotation of responsive genes indicated that SM, WlSM and WgSM differed in the composition and magnitude of transcriptional responses. In SM, the limited set of host-responsive genes was largely restricted to proteolytic and detoxification-associated functions, suggesting targeted fine-tuning rather than broad regulatory remodelling (Supplementary table 3). WlSM additionally modulated regulatory and metabolic genes, consistent with a moderate responsive program. WgSM displayed the broadest response, encompassing genes linked to carbohydrate metabolism, detoxification, and stress-related pathways, with host-dependent directionality. A small conserved host-associated response was shared across morphs: cathepsin B–like proteases were repeatedly up-regulated when aphids fed on oat and ryegrass, irrespective of morph, indicating a shared transcriptional response to specific host nutritional or chemical features. Overall, transcriptional variation is dominated by morph-specific architectures, with host identity influencing a limited set of conserved pathways. We next used co-expression network analyses to dissect the regulatory architecture underlying these strategies.

### Co-expression modules reveal divergent regulatory architectures

To examine coordinated regulation beyond pairwise contrasts, we analyzed co-expression structure across the 1,500 most variable genes and identified six co-expression gene clusters, most structured by morph identity rather than host plant (Fig. 3; Extended data fig. 3). Modules enriched in SM (cluster 2 – 844 genes) were dominated by primary metabolism, detoxification, and cuticle-associated genes and showed little host-dependent modulation. This pattern is consistent with a pre-configured expression landscape. By contrast, several modules reflected regulatory strategies associated with summer morphs and dispersal status. A large cluster shared by both summer morphs (cluster 3; 452 genes) was enriched for chromatin organization and transcriptional regulation, indicating post-transfer reprogramming. WlSM further upregulated ribosome- and translation-associated genes (cluster 1; 113 genes), suggesting a greater reliance on translational/ post-transcriptional control. Notably, SM and WgSM—both retaining migratory or dispersal capacity—exhibited a cluster enriched for stress response, immune-related, and signalling genes, potentially reflecting the physiological demands of migration and transient exposure to novel or suboptimal environments (cluster 4; 70 genes). Host-specific effects were rare: only a small subset of genes exhibited clear morph-by-host interactions (cluster 5; 21), implying that most of the expression variation relevant to acclimation is organized at the morph level. Together, these patterns support canalized/anticipatory regulatory states in SM versus more dynamic post-exposure regulation in summer morphs. We next tested chemical drivers by analyzing host phloem composition.

**Fig. 3:**
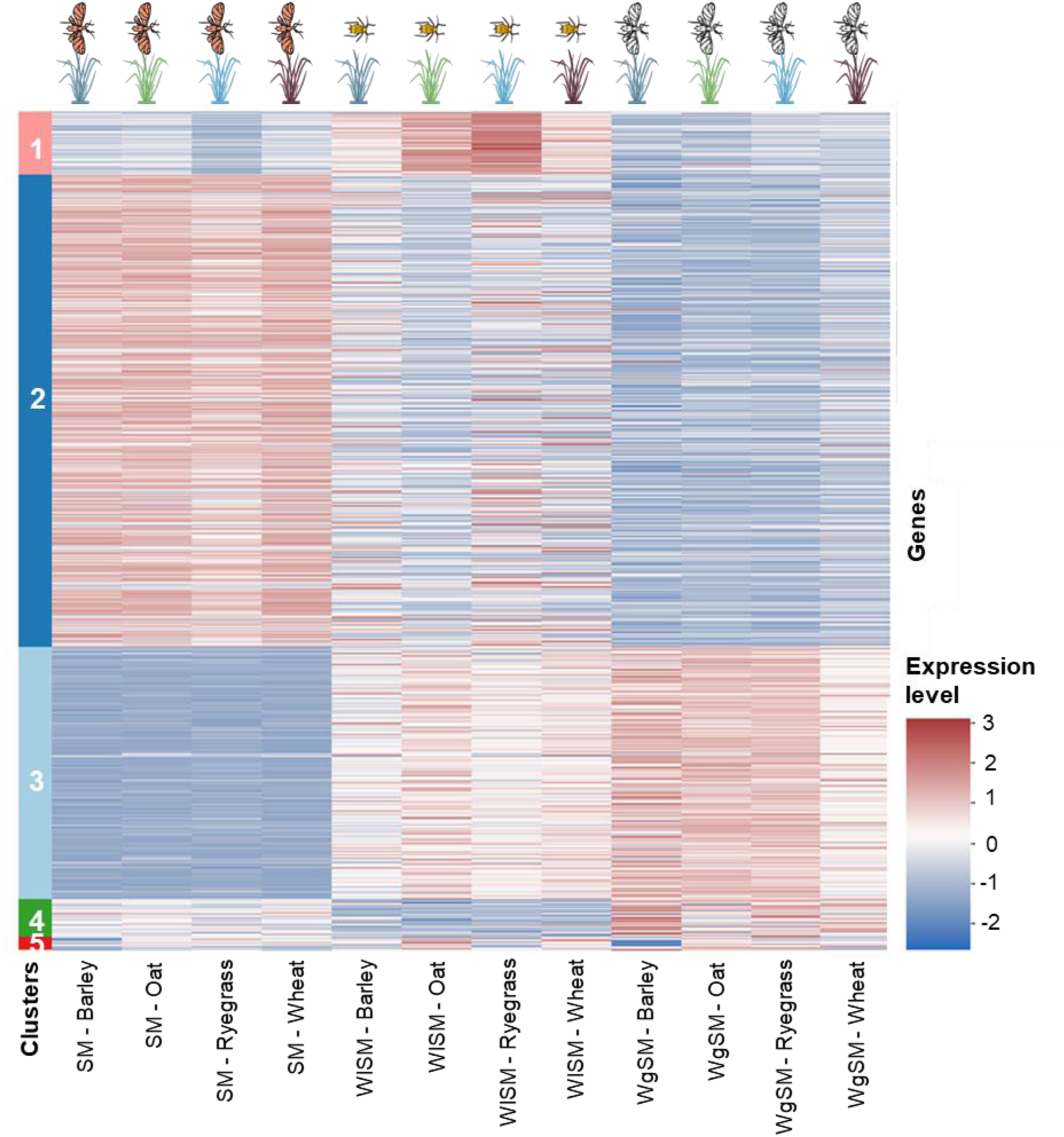
Co-expression heatmap of highly variable genes across morph–host conditions. Heatmap showing z-score–scaled expression profiles of the 1,500 genes with the highest median absolute deviation (MAD), grouped into five co-expression clusters identified by consensus clustering. Colour intensity reflects relative expression levels, with red indicating higher and blue lower expression. Columns correspond to experimental conditions defined by the combination of aphid morphs (SM, WlSM and WgSM) and host plants (barley, oat, wheat and ryegrass), while rows represent individual genes ordered by cluster membership.

### Host plant chemistry shapes nutritional landscapes and reproductive outcomes

To define host nutritional and chemical landscapes, we quantified phloem-associated metabolites across hosts using targeted metabolomics. All Poaceae hosts shared a conserved composition dominated by amino acids, soluble sugars and organic acids (Extended data fig. 4), but differed strongly in the abundance and ratios of these components (Fig. 4; Extended data fig. 5; Supplementary table 4). These quantitative differences generate distinct stoichiometric environments for phloem-feeding aphids (Extended data fig. 6), which must balance high carbon intake, variable nitrogen quality and exposure to defence-associated metabolites while maintaining osmotic homeostasis ^57–59^. Ryegrass, the least suitable host for summer morphs, stood out chemically. It contained the highest total amino acid concentrations (Extended data fig. 6a) but the lowest essential-to-non-essential amino acid ratio (Extended data fig. 6f), indicating nitrogen stored predominantly in forms that require costly conversion for growth. Ryegrass also showed comparatively low sugar-to-amino acid ratios (Extended data fig. 6g), consistent with reduced carbon availability relative to nitrogen pools, and elevated organic acids (Extended data fig. 6c). Among organic acids, quinic acid was strongly enriched in ryegrass compared with the other hosts (Extended data fig. 6e) and emerged as the most discriminating metabolite in multivariate analyses, separating ryegrass from wheat, barley and oat (Fig. 4c). This pattern suggests a link between host defensive allocation (or associated metabolic flux) and aphid performance.

**Fig. 4:**
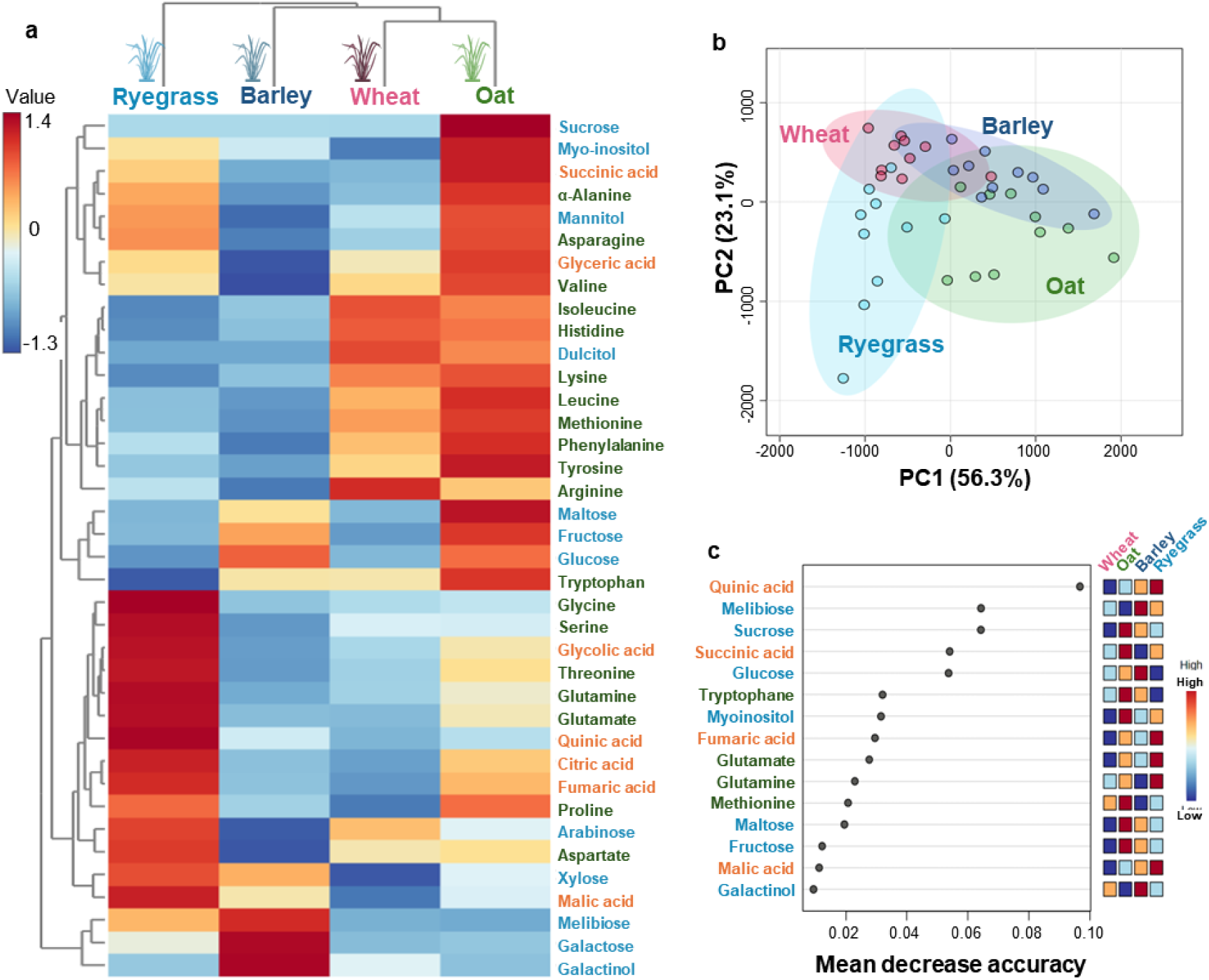
Phloem metabolite profile comparison of the four tested secondary hosts (oat, barley, wheat, and ryegrass) of *Rhopalosiphum padi*. **a,** Hierarchical clustering heatmap of phloem metabolite profiles across the four hosts. Rows represent metabolites and columns represent hosts. Compound font color indicates the chemical class: green = amino acids; orange = organic acids; blue = sugars and sugar alcohols. Asterisks indicate essential amino acids. **b,** Principal component analysis (PCA) scores plot of phloem metabolite profiles from the four host plants. Ellipses represent 95% confidence regions for each host group. **c,** Random-forest variable importance (mean decrease accuracy) ranking phloem metabolites that discriminate the four hosts; right tiles show *z*-scored abundance per host.

To test whether quinic acid contributes causally to reduced host suitability, we experimentally supplemented wheat with quinic acid and measured the performance of wingless summer morphs. Quinic acid supplementation did not significantly affect survival (HR = 20.30, p= 0.99) (Extended data fig. 7a, Supplementary table 5) but caused a pronounced reduction in fecundity relative to acidic water control (estimate = 10.45, *p* < 0.001) (Extended data fig. 7b, Supplementary table 6). Acidic water controls did not reproduce this effect, indicating that reduced reproduction was not attributable to pH alone. These results identify quinic acid as a chemical factor that can directly suppress aphid reproduction in an otherwise suitable host context, providing a mechanistic bridge between host metabolite composition and fitness. In combination with the host metabolomic profiles, the complementation experiment supports the interpretation that the poor performance of summer morphs on ryegrass reflects an integrated constraint involving nutrient balance and defence-associated chemistry, with quinic acid acting as one contributor to reproductive limitation.

## Discussion

This study shows that morphs with distinct ecological roles within a single aphid species deploy contrasting strategies to cope with predictable transitions between unrelated host plants. Spring migrants (SM) maintained uniformly high survival across four chemically distinct grass hosts and exhibited minimal host-induced transcriptional change, whereas summer morphs (WlSM and WgSM) incurred host-dependent survival costs accompanied by broader transcriptional remodeling. Co-expression analyses further revealed that SM are characterized by modules enriched in metabolic and detoxification functions, while summer morphs show stronger involvement of regulatory machinery related to chromatin organization and transcriptional or translational control. Together, these patterns indicate that predictable host alternation can favour a partitioning of plasticity strategies across morphs.

SM appear to rely on a pre-configured physiological state that is broadly compatible with multiple hosts, consistent with anticipatory plasticity evolving under predictable environmental transitions ^11,15,17^. In complex life cycles, such anticipatory states can reduce the fitness costs of abrupt habitat shifts, a central prediction of theory on developmental and life-history plasticity ^3,4,60^. The limited differential expression across the secondary hosts in SM therefore does not reflect weak plasticity; instead, it suggests that the relevant phenotype is already expressed at settlement. This hypothesis is supported by recent work in *R. padi* showing that pre-migratory developmental stages can initiate physiological acclimation prior to host transfer ^42^. This interpretation is consistent with genomic reaction norm theory, which proposes that phenotypic plasticity arises not only from environmentally induced changes in gene expression, but also from baseline gene expression patterns that have been shaped by selection ^61–63^.

By contrast, summer morphs relied on broader transcriptional changes across secondary hosts enriched for chromatin, transcriptional, and translational regulation, consistent with responsive plasticity. Such regulatory architectures reflect the capacity for rapid gene expression reprogramming, increasingly recognized as a central mechanism of insect adaptation to environmental heterogeneity ^37,64^. However, our data show that stronger transcriptional responsiveness did not translate into higher fitness: WgSM exhibited the largest expression changes but the poorest performance on ryegrass. Similar mismatches between transcriptional plasticity and fitness have been reported in other herbivorous insects, suggesting that reactive gene expression may reflect stress or maladaptive responses rather than adaptive acclimation ^65–69^.

Fitness patterns reveal a division of labour between morphs. SM buffered survival across hosts but adjusted fecundity according to host quality, consistent with strong selection to ensure establishment during an irreversible host transition ^42^. Summer morphs, optimized for exploitation of already accepted hosts, experienced survival constraints on chemically challenging hosts, indicating that reactive remodeling may be too slow or too costly to prevent early mortality. Such differences align with broader life-history expectations for aphids, in which SM face predictable but abrupt habitat shifts, whereas resident morphs are optimized for efficient exploitation of accepted hosts ^39,40,43^. This partitioning of plasticity across life-history stages is consistent with theoretical predictions that complex life cycles evolve stage-specific adaptive responses to environmental change, rather than a single uniform strategy ^9,70,71^.

Transcriptomic analysis provides a mechanistic explanation for this life-history contrast between morphs. SM showed limited host-associated differential gene expression, with co-expression modules predominantly associated with metabolic, detoxification, and barrier-related processes involved in gut epithelium, cuticle, and host–plant interaction. In aphids, host alternation is regulated by predictable environmental cues, notably seasonal variation in photoperiod and host-plant physiology ^38,42,72^, which may enable migrants to adopt a broadly compatible physiological state before dispersal. The transcriptional profile observed in SM is therefore consistent with anticipatory plasticity, whereby individuals activate relevant pathways prior to environmental exposure when ecological transitions are predictable ^15,16^. Summer morphs, by contrast, appear to rely mainly on reactive regulation at settlement. Their enrichment for chromatin- and transcription-associated clusters implies stronger upstream regulatory engagement than in SM. This response is consistent with responsive plasticity, in which phenotypes are adjusted following feedback from the current environment, often incurring transient fitness costs during adjustment ^15,73^. Although chromatin and epigenetic mechanisms can support rapid transcriptional change in insects ^37^, this strategy may be less efficient than anticipatory buffering during abrupt but predictable host shifts; for sedentary WlSM, broad pre-activation of metabolic and detoxification programs may be too costly and trade off with fecundity ^43^.

Plant metabolomic analyses further clarify the ecological context in which these contrasting plasticity strategies operate. Host suitability emerged from interactions between nutrient stoichiometry and defence-associated chemistry ^74,75^. Ryegrass combined an unfavourable essential-to-non-essential amino acid and carbon–nitrogen balance with elevated organic acids, conditions known to constrain aphid growth and symbiont-mediated amino acid provisioning ^76–79^. Among these compounds, quinic acid was strongly associated with reduced aphid performance and experimentally shown to suppress fecundity, providing a direct mechanistic link between host chemistry and fitness. Quinic acid and related metabolites have been implicated in plant responses to herbivory and the production of feeding-deterrent compounds ^80–82^. Importantly, these chemical constraints help explain why reactive transcriptional plasticity was insufficient to maintain performance on ryegrass: rather than acting solely through acute toxicity, plant metabolites impose sustained metabolic demands that constrain the ability of aphids to rapidly adjust their physiology. Thus, host chemical landscapes not only determine aphid performance but also shape the selective conditions favoring anticipatory versus responsive plasticity ^34,35^.

Despite strong morph-specific expression landscapes, aphids also exhibited conserved host-associated responses. Notably, cathepsin B–like proteases were consistently upregulated on oat and ryegrass, the two amino acid-abundant hosts, across all morphs, indicating a shared digestive response to hosts with high but stoichiometrically imbalanced nutrient profiles. Cathepsin B proteases are well-established components of aphid digestive physiology and salivary secretions, where they contribute to nutrient acquisition and modulation of plant defences ^34,83,84^. Similarly, detoxification genes such as UDP-glucosyltransferases (UGT) were modulated according to host. Such fine-scale host-associated expression changes involving detoxification enzymes often play a central role in mediating physiological responses to plant chemistry. In herbivorous insects, cytochrome P450 monooxygenases, UGTs, glutathione S-transferases and esterases are widely implicated in the metabolic processing of plant toxins and in the modulation of harmful compounds encountered during feeding ^23,85^. In aphids, cytochrome P450s have been repeatedly linked to host use and acclimation, with expansions and differential expression of P450 gene families associated with adaptation to diverse host plants and insecticide resistance ^34,86^. Such shared responses likely represent a secondary layer of plasticity that fine-tunes digestion and detoxification without overriding morph-specific regulatory architectures.

Our findings provide a mechanistic framework for understanding how phenotypic plasticity evolves in systems with predictable environmental transitions. First, selection can act on the timing and architecture of plastic responses, favoring anticipatory strategies when cues are reliable and responsive strategies when environments are less predictable. Second, plasticity can be partitioned among life-history stages, with different morphs embodying distinct trade-offs between broad tolerance and reproductive efficiency. Third, transcriptomic responsiveness should not be equated with adaptive acclimation: successful performance may depend more on pre-configured expression states than on large transcriptional shifts. More broadly, these results highlight how complex life cycles integrate multiple plasticity strategies within a single genome, linking ecological function to regulatory evolution. By integrating fitness assays, transcriptomics, and metabolomics, we show that morph-specific plasticity strategies emerge from interactions between life-history specialization and host chemical landscapes. Collectively, our study illuminates how phenotypic plasticity evolves in heterogeneous environments and provides a general framework for predicting responses to rapid environmental change.

## Author Contributions

Conceptualization, J-C. Simon, R. Ghosh and A. Etier; methodology, R. Ghosh, A. Etier, M. Berberon, N. Marnet, S. Berardocco; Data analysis, R. Ghosh, A. Etier, M. Berberon, N. Marnet, S. Berardocco; investigation, R. Ghosh, A. Etier; writing—original draft preparation, R. Ghosh, A. Etier; writing—review and editing, J-C. Simon, A. Bouchereau; supervision, J-C. Simon; project administration, J-C. Simon; funding acquisition, J-C. Simon. All authors have read and agreed to the published version of the manuscript.

## Acknowledgments

This work was funded by the European Research Council through the ALTEREVO project (no 101054340) led by J.C. Simon. Bioinformatic analyses have been performed thanks to GenOuest and Genotoul (Toulouse Genopole) servers. We thank Fabrice Legeai and the BIPAA platform for providing access to genome sequence of *R. padi* and bioinformatic tools. We are grateful to Gaëtan Denis, Romuald Cloteau, Yannis Nio, Ségolène Buzy, and Maïwenn Le Floch for their help in fitness experiments. We thank Julie Jaquiery for valuable discussions and advice on statistical analysis.

