## Supplementary files for "Morph-specific plasticity drives divergent host-shift and acclimation strategies in aphids"

### Supplementary figures

**Extended data fig 1. Annual life cycle of *Rhopalosiphum padi* involving host alternation on primary and secondary hosts and experimental design to test morph-specific acclimation to secondary hosts. a,** Schematic representation of the annual life-cycle of *R. padi*. Sexual reproduction and overwintering occur on the woody primary host *Prunus padus* (eggs laid on the primary host). In spring, fundatrices and subsequent parthenogenetic generations produce spring migrants (SM) that emigrate from *P. padus* and colonize grass secondary hosts (Poaceae). During the growing season, populations on grasses consist of wingless summer morphs (WlSM) specialized for local population growth and winged summer morphs (WgSM) that retain dispersal capacity. Winged summer morphs are generally produced in response to population density. In autumn, migrants (gynoparae and males) return to the primary host, where sexual morphs mate and sexual females (oviparae) produce eggs to complete the cycle. **b,** Overview of the host-transfer experiments and multi-omics sampling. Summer morphs were generated on a common secondary host (orchard grass) before transfer. Individuals from three morphs (SM, WlSM, WgSM) were transferred to four secondary hosts (wheat, oat, barley, and ryegrass; Poaceae) to quantify acclimation outcomes. Following transfer, we measured fitness (survival and fecundity) and profiled whole-transcriptome responses in aphids and phloem-associated metabolite composition in host plants (dashed box), enabling integration of performance, regulatory responses, and host chemical landscapes across morphs and hosts.

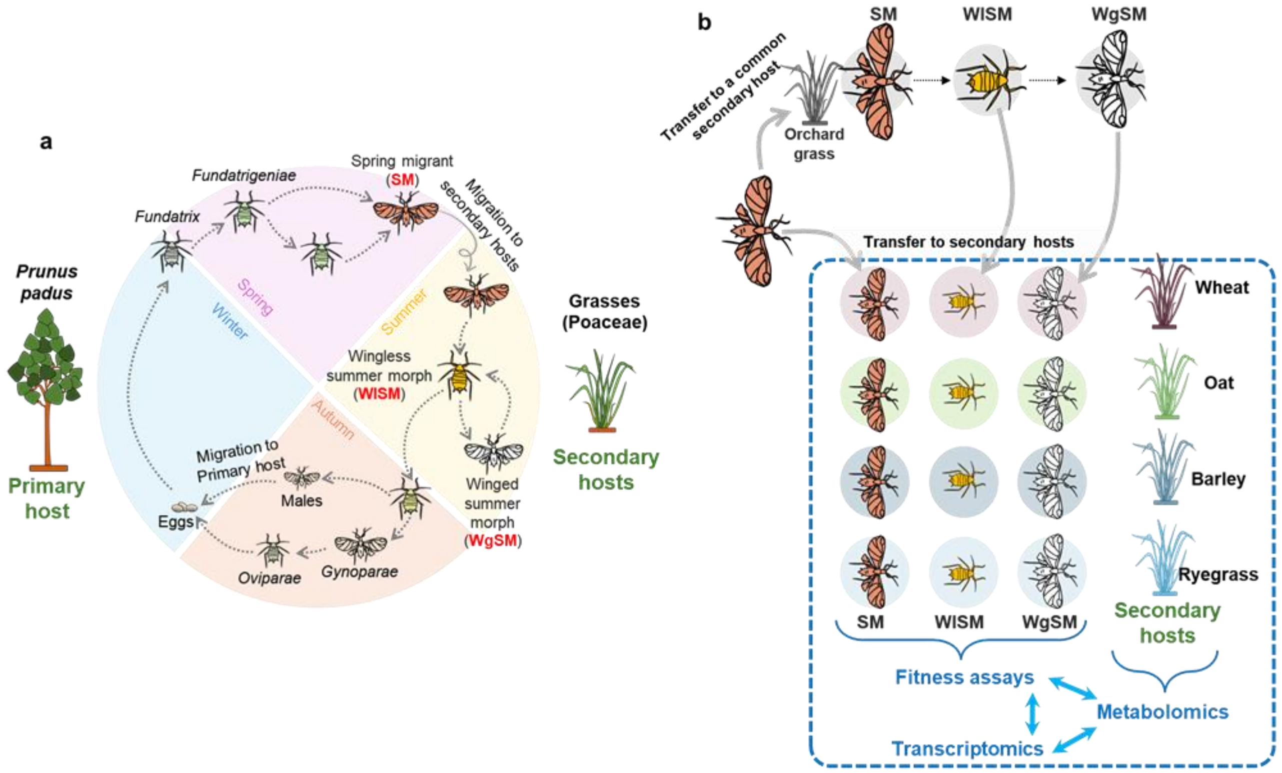

**Extended data fig. 2. Correlation matrix of gene expression profiles of *Rhopalosiphum padi* samples across morphs and host plants.** Correlation plot showing pairwise Pearson correlations among all biological replicates of SM, WlSM, and WgSM reared on wheat, barley, oat, and ryegrass. Replicates are represented by ellipse-shaped symbols whose colour intensity reflects correlation strength (dark green = high correlation; light green = lower correlation). The shape of the ellipses further summarizes correlation structure, with narrow ellipses indicating stronger correlations. Across all morph–host combinations, correlation values ranged from 0.67 to near-perfect correlations, highlighting the overall high transcriptomic similarity among replicates.

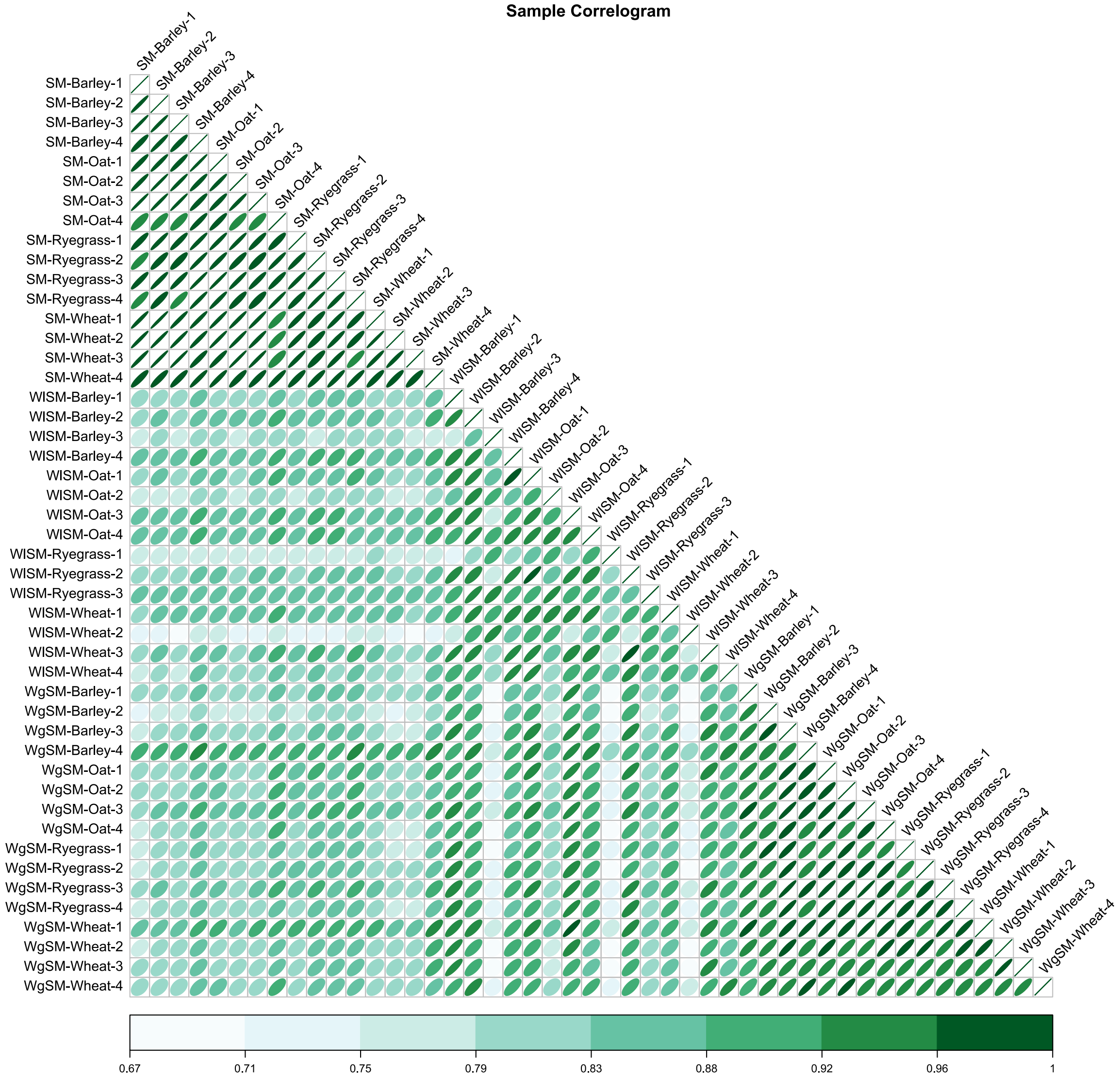

**Extended data fig. 3. Gene Ontology enrichment analysis of co-expression clusters.** Dot plots summarizing enriched Gene Ontology (GO) terms for each co-expression cluster. GO categories are displayed in separate columns corresponding to Biological Process (BP), Cellular Component (CC) and Molecular Function (MF), delineated by dashed lines. Each cluster is framed by a coloured box matching the cluster colour used throughout the study. For each cluster and GO category, the ten most significantly enriched terms are shown, with terms listed on the y-axis and enrichment significance represented on the x-axis as $-\log_{10}$(P-value). Dot size reflects the number of genes associated with each GO term. For clusters or GO categories with no significant enrichment, the label "No GO-term enrichment" is indicated.

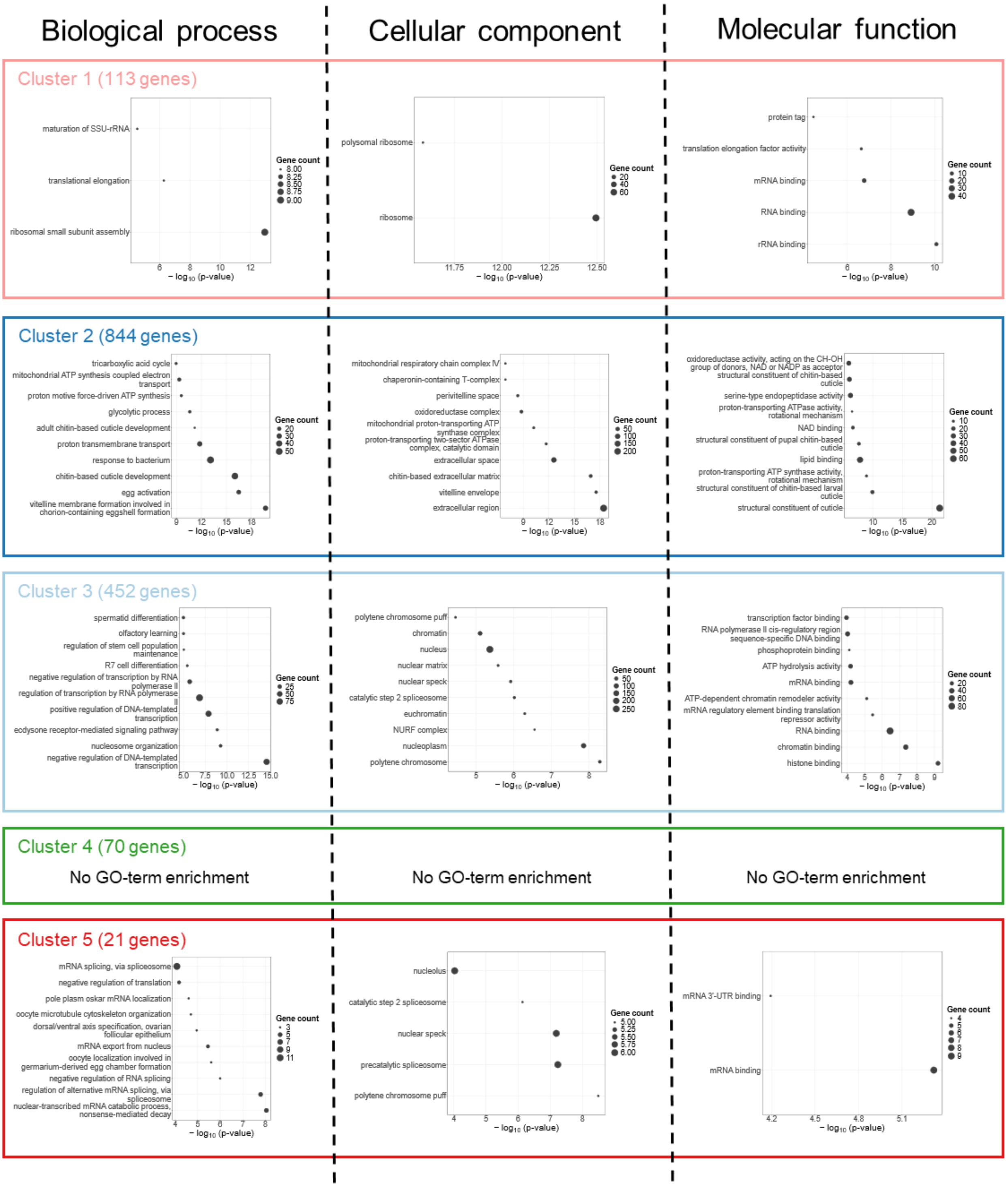

**Extended data fig. 4. Overall phloem metabolite composition across four Poaceae hosts of *Rhopalosiphum padi*. a,** Schematic representation of the phloem collection method. **b,** Boxplots represent the distribution of compound concentrations (nmoles mL$^{-1}$ phloem) pooled across wheat, oat, barley, and ryegrass. Font colors indicate chemical classes: blue– sugars and sugar alcohols, orange– organic acids, green–amino acids. Asterisks indicate essential amino acids.

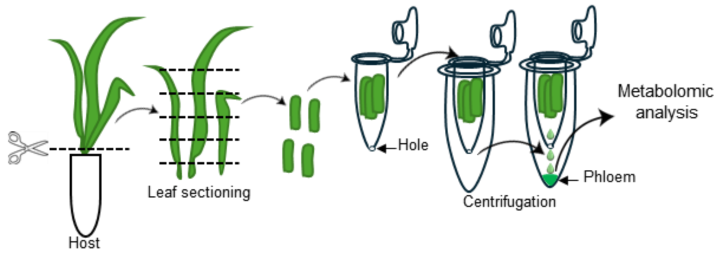


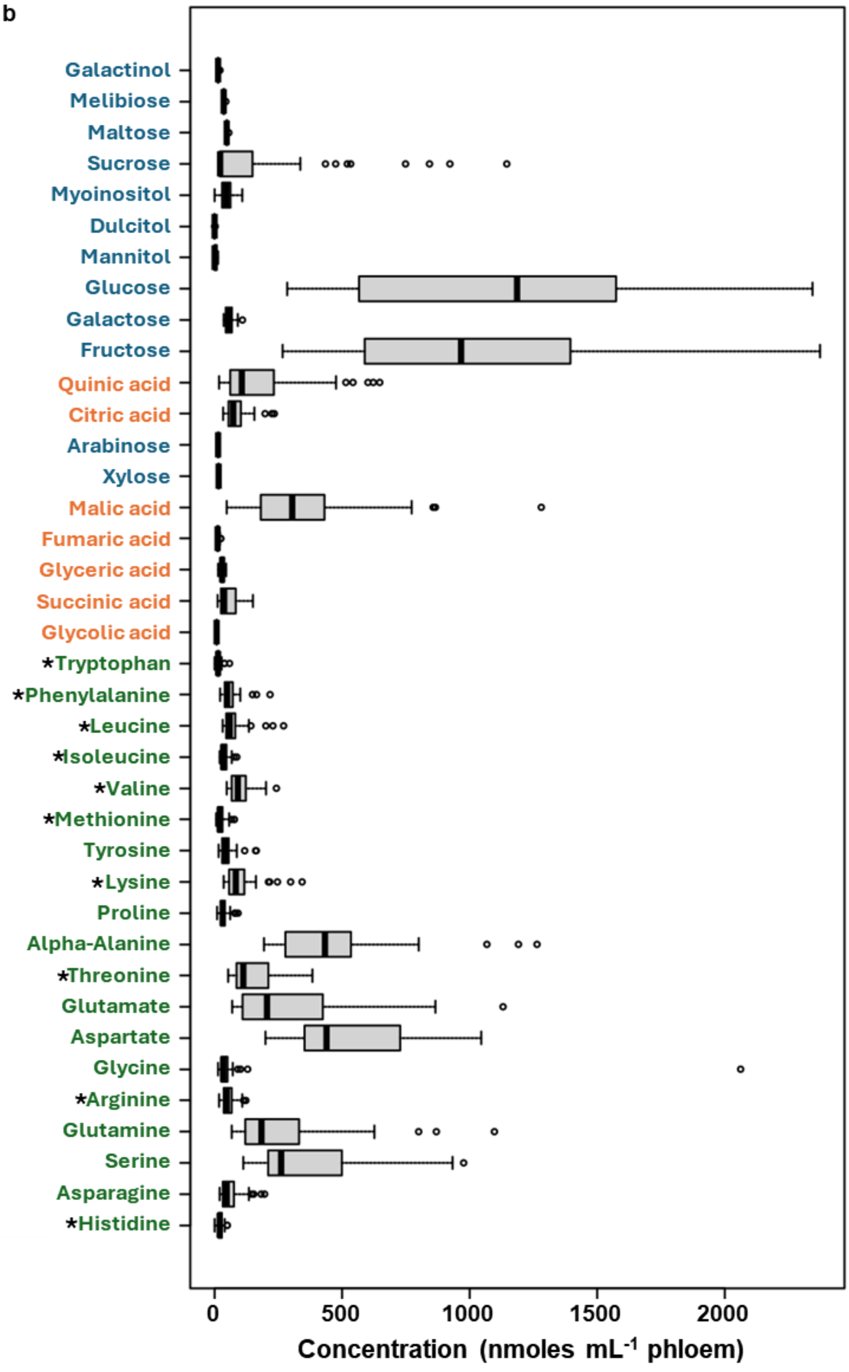

**Extended data fig. 5. Phloem metabolite profile comparison of four secondary hosts of *Rhopalosiphum padi*. a,** The relative proportions of essential amino acids, non-essential amino acids, organic acids, and sugars and sugar alcohols in the phloem sap. **b,** Heatmap depicting phloem metabolite class distribution across the four hosts. Rows represent metabolite classes and columns represent hosts. **c,** The ternary plot represents the relative contributions (%) of amino acids, sugars (including sugar alcohols), and organic acids in four Poaceae hosts. Each point corresponds to a host positioned according to its normalized metabolite composition. The color gradient indicates variation in the nutritional landscape (arbitrary units), highlighting regions of higher versus lower combined nutrient availability.

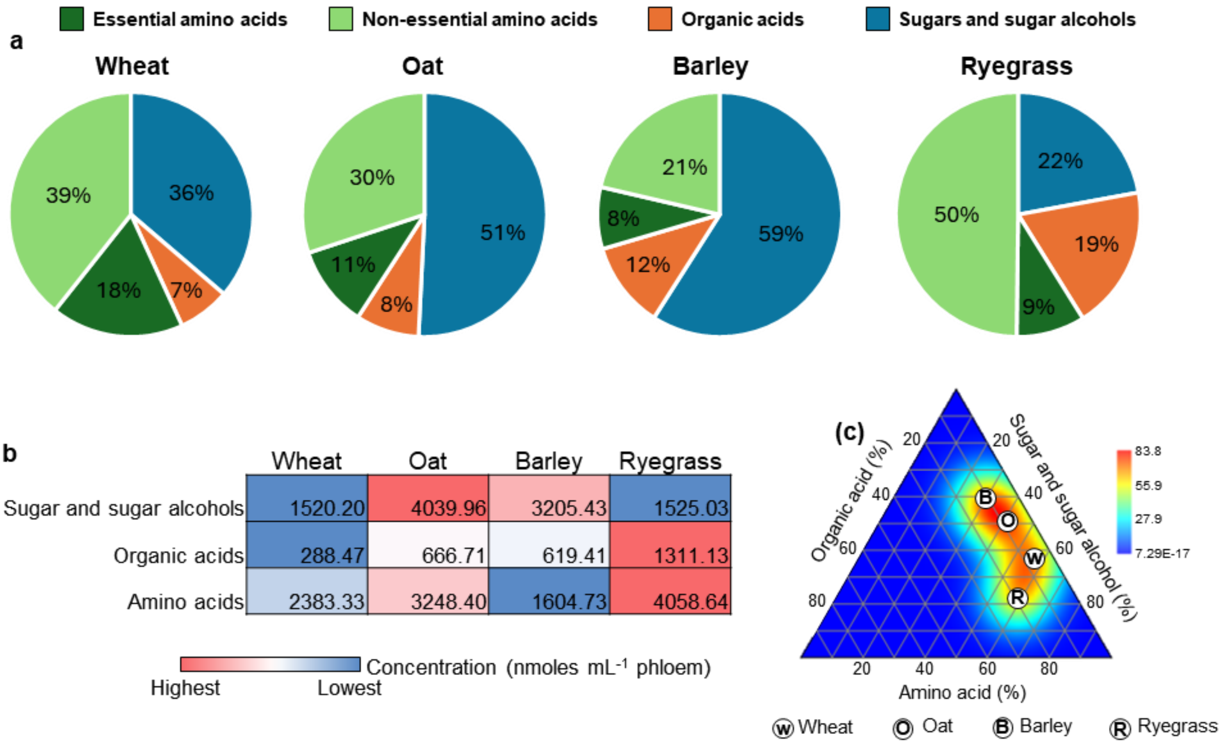

**Extended data fig. 6. Profiles of metabolite concentrations and ratios, illustrating the interspecific variations between four Poaceae hosts of *Rhopalosiphum padi*.** Box plots showing differences in **a,** total amino acid ($\chi^2$ = 18.45), **b,** total sugar (including alcohols) ($\chi^2$ = 27.72), **c,** total organic acid ($\chi^2$ = 29.03), **d,** transport sugar (glucose + fructose + sucrose) ($\chi^2$ = 28.14), **e,** quinic acid ($\chi^2$ = 33.99) concentrations, **f,** essential to nonessential amino acid ($\chi^2$ = 27.68), **g,** sugar to amino acid ($\chi^2$ = 27.68), **h,** sugar to organic acid ($\chi^2$ = 24.68), and **i,** amino acid to organic acid ($\chi^2$ = 17.01) ratios. Different letters above boxes indicate statistically significant differences among host species ($p$ < 0.05), determined using Kruskal–Wallis tests followed by Dunn's post hoc test with Bonferroni corrections.

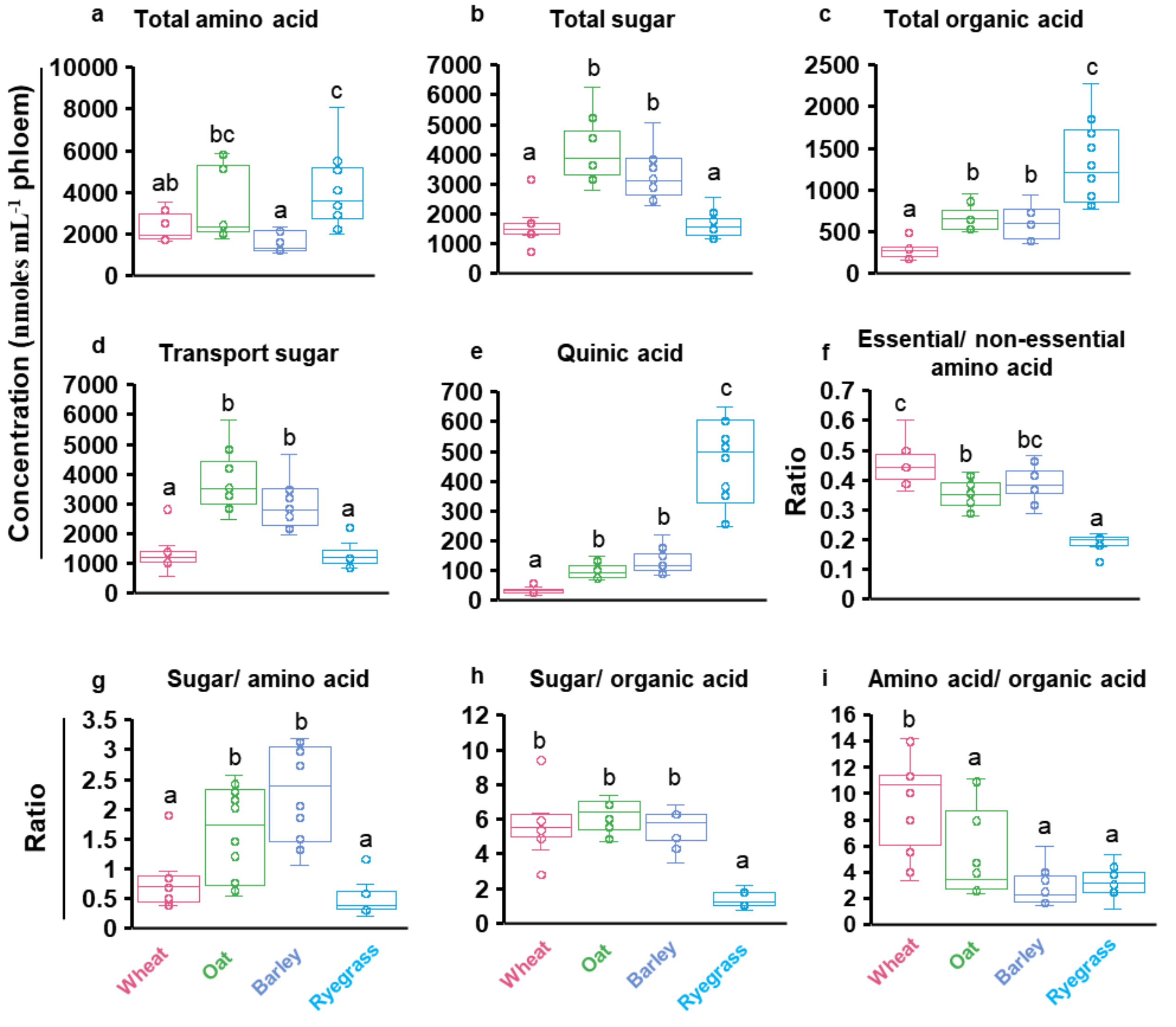

**Extended data fig. 7. Survival and fecundity dynamics of wingless summer morph of *Rhopalosiphum padi* under two chemical conditions: wheat supplemented with acidic water (control) and quinic acid (treatment). a,** show Kaplan–Meier survival curves, and **b,** show cumulative fecundity curves per surviving adult. Each condition was analyzed independently. Shaded areas indicate 95% confidence intervals, and LOESS smoothing was applied to fecundity data for visual clarity.

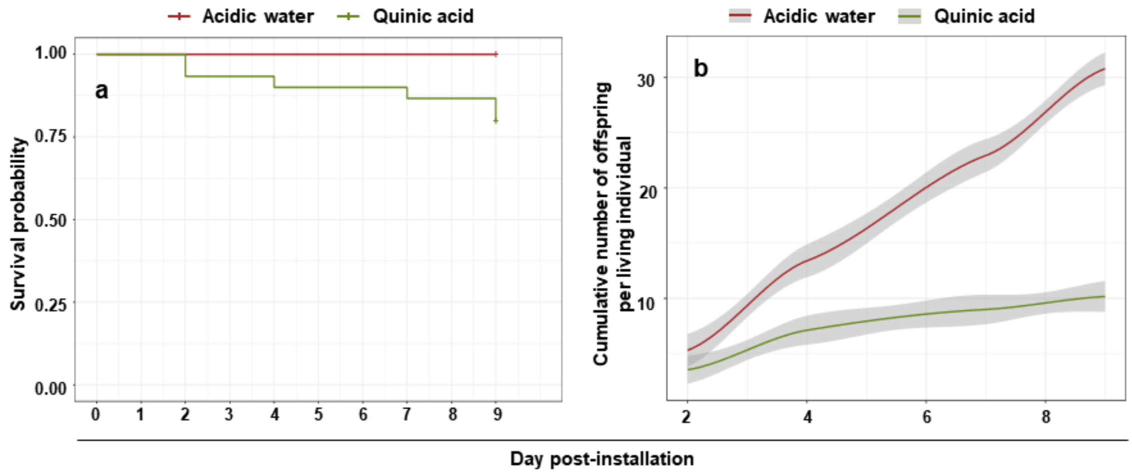

#### Supplementary tables:

**Supplementary table 1. Cox proportional hazards model testing the effect of aphid morph on survival across host plants.** Results of Cox proportional hazards models assessing the association between aphid morph and time to death following transfer onto one of the four host plants. For each pairwise contrast, the regression coefficient (coef), hazard ratio (exp(coef)), standard error (se(coef)), z statistic (z), and associated *P*-value (Pr(>|z|)) are reported. Hazard ratios greater than 1 indicate an increased risk of death relative to the reference level, whereas values below 1 indicate reduced risk.

| Morph | Contrast | coef | exp(coef) | se(coef) | z | Pr(>\|z\|) |
|---|---|---|---|---|---|---|
| SM | Barley vs Oat | 0.092 | 1.096 | 0.535 | 0.172 | 0.863 |
| SM | Barley vs Ryegrass | 0.020 | 1.020 | 0.535 | 0.037 | 0.970 |
| SM | Barley vs Wheat | 0.314 | 1.369 | 0.518 | 0.607 | 0.544 |
| SM | Oat vs Barley | -0.092 | 0.912 | 0.535 | -0.172 | 0.863 |
| SM | Oat vs Ryegrass | -0.072 | 0.930 | 0.535 | -0.135 | 0.893 |
| SM | Oat vs Wheat | 0.222 | 1.249 | 0.518 | 0.430 | 0.668 |
| SM | Ryegrass vs oat | 0.072 | 1.075 | 0.535 | 0.135 | 0.893 |
| SM | Ryegrass vs barley | -0.020 | 0.980 | 0.535 | -0.037 | 0.970 |
| SM | Ryegrass vs Wheat | 0.294 | 1.342 | 0.518 | 0.569 | 0.570 |
| SM | Wheat vs Oat | -0.222 | 0.801 | 0.518 | -0.430 | 0.668 |
| SM | Wheat vs Barley | -0.314 | 0.730 | 0.518 | -0.607 | 0.544 |
| SM | Wheat vs Ryegrass | -0.294 | 0.745 | 0.518 | -0.569 | 0.570 |
| WlSM | Barley vs Oat | -0.191 | 0.826 | 0.379 | -0.503 | 0.615 |
| WlSM | Barley vs Ryegrass | 0.613 | 1.846 | 0.358 | 1.714 | 0.087 |
| WlSM | Barley vs Wheat | -0.314 | 0.731 | 0.387 | -0.810 | 0.418 |
| WlSM | Oat vs Barley | 0.191 | 1.210 | 0.379 | 0.503 | 0.615 |
| WlSM | Oat vs Ryegrass | 0.804 | 2.234 | 0.372 | 2.161 | 0.031 |
| WlSM | Oat vs Wheat | -0.123 | 0.884 | 0.400 | -0.307 | 0.759 |
| WlSM | Ryegrass vs oat | -0.804 | 0.448 | 0.372 | -2.161 | 0.031 |
| WlSM | Ryegrass vs barley | -0.613 | 0.542 | 0.358 | -1.714 | 0.087 |
| WlSM | Ryegrass vs Wheat | -0.927 | 0.396 | 0.381 | -2.430 | 0.015 |
| WlSM | Wheat vs Oat | 0.123 | 1.131 | 0.400 | 0.307 | 0.759 |
| WlSM | Wheat vs Barley | 0.314 | 1.369 | 0.387 | 0.810 | 0.418 |
| WlSM | Wheat vs Ryegrass | 0.927 | 2.527 | 0.381 | 2.430 | 0.015 |
| WgSM | Barley vs Oat | 0.605 | 1.831 | 0.354 | 1.711 | 0.087 |
| WgSM | Barley vs Ryegrass | 0.982 | 2.670 | 0.359 | 2.732 | 0.006 |
| WgSM | Barley vs Wheat | 0.524 | 1.689 | 0.372 | 1.408 | 0.159 |
| WgSM | Oat vs Barley | -0.605 | 0.546 | 0.354 | -1.711 | 0.087 |
| WgSM | Oat vs Ryegrass | 0.377 | 1.458 | 0.330 | 1.141 | 0.254 |
| WgSM | Oat vs Wheat | -0.081 | 0.923 | 0.348 | -0.231 | 0.817 |
| WgSM | Ryegrass vs oat | -0.377 | 0.686 | 0.330 | -1.141 | 0.254 |
| WgSM | Ryegrass vs barley | -0.982 | 0.375 | 0.359 | -2.732 | 0.006 |
| WgSM | Ryegrass vs Wheat | -0.458 | 0.633 | 0.353 | -1.296 | 0.195 |
| WgSM | Wheat vs Oat | 0.081 | 1.084 | 0.348 | 0.231 | 0.817 |
| WgSM | Wheat vs Barley | -0.524 | 0.592 | 0.372 | -1.408 | 0.159 |
| WgSM | Wheat vs Ryegrass | 0.458 | 1.580 | 0.353 | 1.296 | 0.195 |

**Supplementary table 2. Post hoc comparisons of cumulative fecundity among aphid morphs and host plants.** Pairwise contrasts of estimated marginal means derived from linear mixed-effects models testing the effects of morph, host plant, and their interaction on cumulative fecundity. For each contrast, the estimated difference (estimate), standard error (SE), degrees of freedom (df), t statistic (t.ratio), and associated P-value (*p*.value) are reported. P-values were adjusted for multiple testing using Tukey's method. Positive estimates indicate higher fecundity in the first level of the contrast relative to the second.

| Morph | contrast | estimate | SE | df | t.ratio | p.value |
|---|---|---|---|---|---|---|
| SM | Barley vs Oat | -5.350 | 0.824 | 505.118 | -6.492 | 1.38E-09 |
| SM | Barley vs Ryegrass | -0.766 | 0.801 | 502.372 | -0.957 | 0.774 |
| SM | Barley vs Wheat | -5.443 | 0.854 | 504.603 | -6.373 | 2.67E-09 |
| SM | Oat vs Ryegrass | 4.583 | 0.831 | 502.300 | 5.515 | 3.35E-07 |
| SM | Oat vs Wheat | -0.093 | 0.885 | 506.336 | -0.105 | 1.000 |
| SM | Ryegrass vs Wheat | -4.677 | 0.866 | 506.963 | -5.398 | 6.19E-07 |
| WlSM | Barley vs Oat | -1.117 | 1.053 | 506.914 | -1.061 | 0.713 |
| WlSM | Barley vs Ryegrass | 0.234 | 1.381 | 505.782 | 0.169 | 0.998 |
| WlSM | Barley vs Wheat | -0.725 | 1.038 | 506.663 | -0.699 | 0.898 |
| WlSM | Oat vs Ryegrass | 1.351 | 1.365 | 502.117 | 0.990 | 0.755 |
| WlSM | Oat vs Wheat | 0.392 | 1.017 | 503.404 | 0.386 | 0.981 |
| WlSM | Ryegrass vs Wheat | -0.959 | 1.355 | 501.736 | -0.707 | 0.894 |
| WgSM | Barley vs Oat | 1.677 | 0.953 | 504.234 | 1.758 | 0.295 |
| WgSM | Barley vs Wheat | -0.864 | 1.058 | 506.957 | -0.816 | 0.847 |
| WgSM | Oat vs Wheat | -2.541 | 1.118 | 506.914 | -2.273 | 0.106 |

**Supplementary table 3. Host-associated differential expression of genes between host-plant contrasts of e genes in the SM, WlSM and WgSM morphs.** For each row, the contrast indicates the pairwise host comparison in which the gene is differentially expressed, *Gene_id* corresponds to the *R. padi* gene identifier, *logFC* represents the $\log_2$ fold change in expression between hosts, and *Function* reports the functional annotation of the gene.

| Contrast | Gene_id | logFC | Function |
|---|---|---|---|
| SM-Barley vs SM-Oat | RPAD4_8688 | -2,521502298 | cathepsin B-like, Expressed effectors |
| SM-Barley vs SM-Oat | RPAD4_5216 | -1,775861639 | cathepsin B-like |
| SM-Barley vs SM-Oat | RPAD4_6460 | -1,017596984 | UDP-glucosyltransferase 2-like |
| SM-Barley vs SM-Oat | RPAD4_12183 | -2,953364279 | zinc finger MYM-type protein 1-like |
| SM-Barley vs SM-Oat | RPAD4_14652 | -1,213503559 | putative 3-methyladenine DNA glycosylase |
| SM-Oat vs SM-Wheat | RPAD4_8688 | 2,850480414 | cathepsin B-like, Expressed effectors |
| SM-Oat vs SM-Wheat | RPAD4_5216 | 1,826061052 | cathepsin B-like |
| SM-Oat vs SM-Wheat | RPAD4_13033 | 1,094032325 | uncharacterized LOC132929882 |
| SM-Ryegrass vs SM-Wheat | RPAD4_8688 | 2,858194018 | cathepsin B-like, Expressed effectors |
| SM-Ryegrass vs SM-Wheat | RPAD4_13033 | 1,582090702 | uncharacterized LOC132929882 |
| SM-Ryegrass vs SM-Wheat | RPAD4_8434 | 1,170120288 | 52 kDa repressor of the inhibitor of the protein kinase-like, Up effectors, Expressed effectors |
| SM-Barley vs SM-Ryegrass | RPAD4_8688 | -2,529215901 | cathepsin B-like, Expressed effectors |
| SM-Barley vs SM-Ryegrass | RPAD4_11564 | 2,220957008 | uncharacterized LOC132925574 |
| SM-Barley vs SM-Ryegrass | RPAD4_6129 | -2,112453871 | UDP-glucosyltransferase 2-like |

| Contrast | Gene_id | logFC | Function |
|---|---|---|---|
| SM-Barley vs SM-Ryegrass | RPAD4_12183 | -3,91225014 | zinc finger MYM-type protein 1-like |
| SM-Barley vs SM-Ryegrass | RPAD4_9977 | -1,587471537 | uncharacterized LOC132927873 |
| SM-Barley vs SM-Wheat | RPAD4_6129 | -1,697857312 | UDP-glucosyltransferase 2-like |
| WlSM-Barley vs WlSM-Ryegrass | RPAD4_8688 | -6,160162906 | cathepsin B-like, Expressed effectors |
| WlSM-Barley vs WlSM-Ryegrass | RPAD4_14298 | -2,021627818 | uncharacterized LOC132930385 |
| WlSM-Barley vs WlSM-Ryegrass | RPAD4_8687 | -5,99832959 | cathepsin B-like, Expressed effectors |
| WlSM-Barley vs WlSM-Ryegrass | RPAD4_4763 | -1,839872836 | histone H1A, sperm-like |
| WlSM-Barley vs WlSM-Ryegrass | RPAD4_11802 | 10,14229522 | uncharacterized LOC132923759 |
| WlSM-Oat vs WlSM-Barley | RPAD4_8688 | 4,627319024 | cathepsin B-like, Expressed effectors |
| WlSM-Oat vs WlSM-Barley | RPAD4_1683 | 2,079644467 | uncharacterized LOC132918407 |
| WlSM-Oat vs WlSM-Barley | RPAD4_770 | -1,813156415 | lipase 3-like, Expressed effectors |
| WlSM-Oat vs WlSM-Barley | RPAD4_7948 | -1,531759332 | myogenesis-regulating glycosidase-like, Expressed effectors |
| WlSM-Oat vs WlSM-Barley | RPAD4_8687 | 4,566184489 | cathepsin B-like, Expressed effectors |
| WlSM-Oat vs WlSM-Barley | RPAD4_5066 | -1,077551873 | delta(24)-sterol reductase-like |
| WlSM-Oat vs WlSM-Barley | RPAD4_1688 | 1,081199923 | nuclear receptor ROR-alpha B-like |
| WlSM-Oat vs WlSM-Barley | RPAD4_1889 | -1,262914034 | cytochrome P450 4C1-like |
| WlSM-Oat vs WlSM-Barley | RPAD4_3782 | 1,096922576 | G patch domain-containing protein 4 |
| WlSM-Oat vs WlSM-Barley | RPAD4_3978 | 1,573975374 | putative uncharacterized protein DDB_G0271982 |
| WlSM-Oat vs WlSM-Barley | RPAD4_754 | 1,271203594 | uncharacterized LOC132931762 |
| WlSM-Oat vs WlSM-Barley | RPAD4_4837 | 1,019239805 | histone H1A, sperm-like |
| WlSM-Oat vs WlSM-Barley | RPAD4_5278 | 1,133658251 | receptor expression-enhancing protein 5-like |
| WlSM-Oat vs WlSM-Barley | RPAD4_6129 | 1,860555789 | UDP-glucosyltransferase 2-like |
| WlSM-Oat vs WlSM-Barley | RPAD4_6876 | 1,181068895 | uncharacterized LOC132920782 |
| WlSM-Oat vs WlSM-Barley | RPAD4_10673 | 1,237464896 | uncharacterized LOC132926772 |
| WlSM-Oat vs WlSM-Barley | RPAD4_10865 | 1,080055684 | uncharacterized LOC132924328 |
| WlSM-Oat vs WlSM-Barley | RPAD4_11214 | 1,078544393 | uncharacterized LOC132925373 |
| WlSM-Oat vs WlSM-Barley | RPAD4_11878 | 1,149185768 | uncharacterized LOC132926328 |
| WlSM-Oat vs WlSM-Barley | RPAD4_12532 | 1,033521548 | cytochrome P450 4C1-like |
| WlSM-Oat vs WlSM-Barley | RPAD4_13318 | -1,511418234 | cuticlin-4-like |
| WlSM-Oat vs WlSM-Barley | RPAD4_15045 | 1,143768279 | calphotin-like |
| WlSM-Oat vs WlSM-Barley | RPAD4_7856 | -1,050019112 | basic proline-rich protein-like |
| WlSM-Oat vs WlSM-Barley | RPAD4_4758 | -3,554287333 | histone H2A |
| WlSM-Oat vs WlSM-Barley | RPAD4_9200 | -1,254281 | uncharacterized LOC132927203 |
| WlSM-Oat vs WlSM-Barley | RPAD4_12140 | 2,097160481 | uncharacterized LOC132927527 |
| WlSM-Oat vs WlSM-Barley | RPAD4_9223 | -1,747842015 | uncharacterized LOC132924053 |
| WlSM-Oat vs WlSM-Wheat | RPAD4_8688 | 5,135200016 | cathepsin B-like, Expressed effectors |
| WlSM-Oat vs WlSM-Wheat | RPAD4_8687 | 6,314821081 | cathepsin B-like, Expressed effectors |
| WlSM-Oat vs WlSM-Wheat | RPAD4_770 | -2,137811362 | lipase 3-like, Expressed effectors |
| WlSM-Oat vs WlSM-Wheat | RPAD4_1683 | 1,739249013 | uncharacterized LOC132918407 |
| WlSM-Oat vs WlSM-Wheat | RPAD4_9436 | 1,165611367 | probable phospholipid-transporting ATPase IA |
| WlSM-Oat vs WlSM-Wheat | RPAD4_6552 | -1,379489648 | uncharacterized LOC132919007 |
| WlSM-Oat vs WlSM-Wheat | RPAD4_13475 | 1,221570042 | plexin-A2-like |
| WlSM-Oat vs WlSM-Wheat | RPAD4_4606 | -2,847189473 | cuticle protein 7-like |

| Contrast | Gene_id | logFC | Function |
|---|---|---|---|
| WlSM-Oat vs WlSM-Wheat | RPAD4_105 | -2,265134866 | uncharacterized LOC132918644 |
| WlSM-Oat vs WlSM-Wheat | RPAD4_1688 | 1,009532088 | nuclear receptor ROR-alpha B-like |
| WlSM-Oat vs WlSM-Wheat | RPAD4_731 | -1,159742585 | bromodomain-containing protein DDB_G0280777-like |
| WlSM-Oat vs WlSM-Wheat | RPAD4_6308 | -3,032505021 | uncharacterized LOC132920294 |
| WlSM-Oat vs WlSM-Wheat | RPAD4_7199 | 1,027264422 | G-protein coupled receptor dmsr-1-like |
| WlSM-Oat vs WlSM-Wheat | RPAD4_7947 | -1,750196937 | myogenesis-regulating glycosidase-like, Expressed effectors |
| WlSM-Oat vs WlSM-Wheat | RPAD4_10259 | -1,262416622 | uncharacterized LOC132924465 |
| WlSM-Oat vs WlSM-Wheat | RPAD4_13838 | -1,127565746 | thw - chitin-binding domain protein thawb, Expressed effectors |
| WlSM-Wheat vs WlSM-Barley | RPAD4_6129 | 4,31291489 | UDP-glucosyltransferase 2-like |
| WlSM-Wheat vs WlSM-Barley | RPAD4_1174 | -1,122475509 | proton channel OtopLc-like |
| WlSM-Wheat vs WlSM-Barley | RPAD4_5220 | -1,447876152 | uncharacterized LOC132923177 |
| WlSM-Wheat vs WlSM-Barley | RPAD4_7056 | -1,126595362 | uncharacterized LOC132920639 |
| WlSM-Wheat vs WlSM-Barley | RPAD4_11616 | 10,16968094 | uncharacterized LOC132927064 |
| WlSM-Wheat vs WlSM-Barley | RPAD4_8952 | -1,287043303 | uncharacterized LOC132927184 |
| WlSM-Wheat vs WlSM-Barley | RPAD4_9223 | -1,173280072 | uncharacterized LOC132924053 |
| WlSM-Wheat vs WlSM-Barley | RPAD4_9544 | -1,112773092 | uncharacterized LOC132927463 |
| WlSM-Wheat vs WlSM-Barley | RPAD4_13247 | -1,044958226 | neuroendocrine convertase 1-like |
| WlSM-Wheat vs WlSM-Ryegrass | RPAD4_8688 | -6,668043898 | cathepsin B-like, Expressed effectors |
| WlSM-Wheat vs WlSM-Ryegrass | RPAD4_8687 | -7,746966182 | cathepsin B-like, Expressed effectors |
| WlSM-Wheat vs WlSM-Ryegrass | RPAD4_14298 | -2,370488212 | uncharacterized LOC132930385 |
| WlSM-Wheat vs WlSM-Ryegrass | RPAD4_14904 | -1,751897247 | elongation of very long chain fatty acids protein 4-like |
| WlSM-Wheat vs WlSM-Ryegrass | RPAD4_12972 | -1,381019543 | uncharacterized LOC132928716 |
| WgSM-Wheat vs WgSM-Ryegrass | RPAD4_8688 | -5,191631055 | cathepsin B-like, Expressed effectors |
| WgSM-Wheat vs WgSM-Ryegrass | RPAD4_8687 | -5,91134156 | cathepsin B-like, Expressed effectors |
| WgSM-Wheat vs WgSM-Ryegrass | RPAD4_12628 | 1,301125207 | uncharacterized LOC132929792 |
| WgSM-Wheat vs WgSM-Ryegrass | RPAD4_7192 | 1,712122546 | uncharacterized LOC132921431 |
| WgSM-Wheat vs WgSM-Ryegrass | RPAD4_14465 | 1,362750434 | uncharacterized LOC132930328 |
| WgSM-Wheat vs WgSM-Ryegrass | RPAD4_14197 | 1,329387115 | maltase 2-like |
| WgSM-Wheat vs WgSM-Ryegrass | RPAD4_5776 | 1,330921044 | uncharacterized LOC132920535 |
| WgSM-Wheat vs WgSM-Ryegrass | RPAD4_10014 | 1,54478331 | uncharacterized LOC132927283 |
| WgSM-Wheat vs WgSM-Ryegrass | RPAD4_13072 | 1,082735534 | UDP-glucosyltransferase 2-like |
| WgSM-Wheat vs WgSM-Ryegrass | RPAD4_5843 | 1,907891187 | cytochrome P450 6k1-like |
| WgSM-Wheat vs WgSM-Ryegrass | RPAD4_1327 | 1,461524614 | uncharacterized LOC132917367 |
| WgSM-Wheat vs WgSM-Ryegrass | RPAD4_15158 | 1,020491453 | sel - canopy family protein seele, Expressed effectors |
| WgSM-Wheat vs WgSM-Ryegrass | RPAD4_5990 | 1,981673332 | phenoloxidase-activating factor 2-like |
| WgSM-Wheat vs WgSM-Ryegrass | RPAD4_14464 | 1,163365922 | uncharacterized LOC132930276 |

| Contrast | Gene_id | logFC | Function |
|---|---|---|---|
| WgSM-Wheat vs WgSM-Ryegrass | RPAD4_15118 | 1,094971561 | uncharacterized LOC132928306 |
| WgSM-Wheat vs WgSM-Ryegrass | RPAD4_945 | 1,127685318 | uncharacterized LOC132917697 |
| WgSM-Wheat vs WgSM-Ryegrass | RPAD4_13025 | -1,350348793 | glycerol kinase-like |
| WgSM-Wheat vs WgSM-Ryegrass | RPAD4_7825 | 1,436805221 | small integral membrane protein 8 |
| WgSM-Wheat vs WgSM-Ryegrass | RPAD4_1312 | 1,112038326 | Karl - lipocalin/cytosolic fatty acid-binding protein Karl, Expressed effectors |
| WgSM-Wheat vs WgSM-Ryegrass | RPAD4_14649 | 1,12654198 | germ cell nuclear acidic protein-like |
| WgSM-Wheat vs WgSM-Ryegrass | RPAD4_379 | -1,151266012 | protein maelstrom 1-like |
| WgSM-Wheat vs WgSM-Ryegrass | RPAD4_749 | 1,08177674 | cathepsin B-like cysteine proteinase 4, Expressed effectors |
| WgSM-Wheat vs WgSM-Ryegrass | RPAD4_6689 | 1,542786048 | uncharacterized LOC132922845 |
| WgSM-Wheat vs WgSM-Ryegrass | RPAD4_10858 | 1,162431271 | uncharacterized LOC132924179 |
| WgSM-Wheat vs WgSM-Ryegrass | RPAD4_9522 | -1,481675863 | dynein axonemal heavy chain 3-like |
| WgSM-Wheat vs WgSM-Ryegrass | RPAD4_9604 | 1,186119811 | putative fatty acyl-CoA reductase CG5065 |
| WgSM-Wheat vs WgSM-Ryegrass | RPAD4_12605 | 1,214944202 | mucin-2-like |
| WgSM-Wheat vs WgSM-Ryegrass | RPAD4_3495 | 1,029805352 | Ptr - patched domain-containing protein |
| WgSM-Wheat vs WgSM-Ryegrass | RPAD4_13391 | 1,038629175 | 4-hydroxy-2-oxoglutarate aldolase, mitochondrial-like |
| WgSM-Wheat vs WgSM-Ryegrass | RPAD4_4805 | 1,323221199 | nose resistant to fluoxetine protein 6-like |
| WgSM-Wheat vs WgSM-Ryegrass | RPAD4_7856 | 1,752652757 | basic proline-rich protein-like |
| WgSM-Wheat vs WgSM-Ryegrass | RPAD4_10013 | 1,122921604 | uncharacterized LOC132927138 |
| WgSM-Wheat vs WgSM-Ryegrass | RPAD4_13009 | 1,225975961 | probable cytochrome P450 6a13 |
| WgSM-Wheat vs WgSM-Ryegrass | RPAD4_7683 | 1,164551651 | uncharacterized LOC132922088 |
| WgSM-Wheat vs WgSM-Ryegrass | RPAD4_5369 | 1,503466454 | scoloptoxin SSD14-like |
| WgSM-Wheat vs WgSM-Ryegrass | RPAD4_1377 | 1,396185553 | uncharacterized LOC132932166 |
| WgSM-Wheat vs WgSM-Ryegrass | RPAD4_6945 | 1,28758036 | uncharacterized LOC132921646 |
| WgSM-Wheat vs WgSM-Ryegrass | RPAD4_14298 | -1,099251464 | uncharacterized LOC132930385 |
| WgSM-Wheat vs WgSM-Ryegrass | RPAD4_4461 | 1,004118541 | proton-coupled amino acid transporter 2-like |
| WgSM-Wheat vs WgSM-Ryegrass | RPAD4_3799 | -1,056024657 | uncharacterized LOC132917972 |
| WgSM-Wheat vs WgSM-Ryegrass | RPAD4_3899 | -1,388690801 | cathepsin B-like |
| WgSM-Wheat vs WgSM-Ryegrass | RPAD4_15117 | 1,05048902 | prisilkin-39-like, Up effectors, Expressed effectors |
| WgSM-Wheat vs WgSM-Ryegrass | RPAD4_967 | 1,533707537 | acetyl-coenzyme A transporter 1-like |
| WgSM-Wheat vs WgSM-Ryegrass | RPAD4_3837 | 1,070618314 | uncharacterized LOC132917960 |

| Contrast | Gene_id | logFC | Function |
|---|---|---|---|
| WgSM-Wheat vs WgSM-Ryegrass | RPAD4_13247 | 1,417468907 | neuroendocrine convertase 1-like |
| WgSM-Wheat vs WgSM-Ryegrass | RPAD4_7947 | 1,242580689 | myogenesis-regulating glycosidase-like, Expressed effectors |
| WgSM-Wheat vs WgSM-Ryegrass | RPAD4_131 | 1,246316547 | elongation of very long chain fatty acids protein 7-like |
| WgSM-Wheat vs WgSM-Ryegrass | RPAD4_1090 | 1,023284302 | UDP-glucosyltransferase 2-like |
| WgSM-Wheat vs WgSM-Ryegrass | RPAD4_11631 | -1,179834692 | NF-kappa-B-activating protein-like |
| WgSM-Wheat vs WgSM-Ryegrass | RPAD4_4799 | 1,010628209 | fatty acid synthase-like |
| WgSM-Wheat vs WgSM-Ryegrass | RPAD4_200 | 3,342689915 | uncharacterized LOC132918685 |
| WgSM-Wheat vs WgSM-Ryegrass | RPAD4_2449 | 1,076016068 | sepiapterin reductase-like |
| WgSM-Wheat vs WgSM-Ryegrass | RPAD4_8172 | 1,001381575 | probable cytochrome P450 6a13 |
| WgSM-Wheat vs WgSM-Ryegrass | RPAD4_15120 | 1,395839273 | AT-rich interactive domain-containing protein 1A-like |
| WgSM-Wheat vs WgSM-Ryegrass | RPAD4_15135 | 1,043653418 | protein D3-like |
| WgSM-Barley vs WgSM-Ryegrass | RPAD4_8688 | -5,281607874 | cathepsin B-like, Expressed effectors |
| WgSM-Barley vs WgSM-Ryegrass | RPAD4_13718 | 2,509388858 | glucose dehydrogenase [FAD, quinone]-like |
| WgSM-Barley vs WgSM-Ryegrass | RPAD4_2545 | 1,697695854 | dynein axonemal heavy chain 7-like |
| WgSM-Barley vs WgSM-Ryegrass | RPAD4_10107 | 1,38781114 | Dhc93AB - Dynein heavy chain at 93AB |
| WgSM-Barley vs WgSM-Ryegrass | RPAD4_13530 | 1,267190006 | uncharacterized LOC132930100 |
| WgSM-Barley vs WgSM-Ryegrass | RPAD4_14298 | -1,549068508 | uncharacterized LOC132930385 |
| WgSM-Barley vs WgSM-Ryegrass | RPAD4_14758 | 1,059063742 | myosin-J heavy chain |
| WgSM-Barley vs WgSM-Ryegrass | RPAD4_13529 | 3,058040185 | uncharacterized LOC132930099 |
| WgSM-Barley vs WgSM-Ryegrass | RPAD4_1931 | 1,001376284 | uncharacterized LOC132918277 |
| WgSM-Barley vs WgSM-Ryegrass | RPAD4_8687 | -3,308293231 | cathepsin B-like, Expressed effectors |
| WgSM-Barley vs WgSM-Ryegrass | RPAD4_6774 | 1,437842031 | solute carrier family 2, facilitated glucose transporter member 8-like |
| WgSM-Barley vs WgSM-Ryegrass | RPAD4_8217 | -1,570564559 | uncharacterized LOC132922903 |
| WgSM-Barley vs WgSM-Ryegrass | RPAD4_3899 | -1,557287218 | cathepsin B-like |
| WgSM-Barley vs WgSM-Ryegrass | RPAD4_12194 | 1,007879796 | uncharacterized LOC132925991 |
| WgSM-Barley vs WgSM-Ryegrass | RPAD4_14303 | 1,010266112 | hexokinase-4-like |
| WgSM-Barley vs WgSM-Ryegrass | RPAD4_6869 | 1,318594253 | uncharacterized LOC132922618 |
| WgSM-Barley vs WgSM-Ryegrass | RPAD4_5990 | 1,510431659 | phenoloxidase-activating factor 2-like |
| WgSM-Barley vs WgSM-Ryegrass | RPAD4_6868 | 1,104254705 | uncharacterized LOC132922617 |
| WgSM-Barley vs WgSM-Ryegrass | RPAD4_7198 | 1,235141421 | zinc finger protein 771-like |

| Contrast | Gene_id | logFC | Function |
|---|---|---|---|
| WgSM-Barley vs WgSM-Ryegrass | RPAD4_15135 | 1,484662955 | protein D3-like |
| WgSM-Barley vs WgSM-Ryegrass | RPAD4_3476 | -1,245469399 | polyprenol reductase |
| WgSM-Barley vs WgSM-Ryegrass | RPAD4_566 | 1,337159728 | Osi20 - DUF1676 domain-containing protein Osi20 |
| WgSM-Barley vs WgSM-Ryegrass | RPAD4_5712 | 1,534170828 | uncharacterized LOC132922583 |
| WgSM-Barley vs WgSM-Ryegrass | RPAD4_8002 | 1,058493463 | uncharacterized LOC132920223 |
| WgSM-Barley vs WgSM-Ryegrass | RPAD4_8172 | 1,267389021 | probable cytochrome P450 6a13 |
| WgSM-Barley vs WgSM-Ryegrass | RPAD4_9300 | 1,298886044 | uncharacterized LOC132927049 |
| WgSM-Barley vs WgSM-Ryegrass | RPAD4_9532 | 1,266770288 | uncharacterized LOC132927254 |
| WgSM-Barley vs WgSM-Ryegrass | RPAD4_15117 | 1,051822145 | prisilkin-39-like, Up effectors, Expressed effectors |
| WgSM-Barley vs WgSM-Ryegrass | RPAD4_15119 | 1,036392663 | uncharacterized LOC132928307 |
| WgSM-Barley vs WgSM-Ryegrass | RPAD4_9247 | 1,535298862 | dynein axonemal intermediate chain 4-like |
| WgSM-Barley vs WgSM-Ryegrass | RPAD4_5988 | 1,292062689 | uncharacterized LOC132922471 |
| WgSM-Barley vs WgSM-Ryegrass | RPAD4_12809 | 2,205022386 | TM2 domain-containing protein CG11103-like |
| WgSM-Barley vs WgSM-Ryegrass | RPAD4_6037 | -3,650200191 | sugar transporter SWEET1-like |
| WgSM-Oat vs WgSM-Barley | RPAD4_8688 | 3,732370709 | cathepsin B-like, Expressed effectors |
| WgSM-Oat vs WgSM-Barley | RPAD4_9593 | 1,588742403 | facilitated trehalose transporter Tret1-like |
| WgSM-Oat vs WgSM-Barley | RPAD4_13530 | -1,281796101 | uncharacterized LOC132930100 |
| WgSM-Oat vs WgSM-Barley | RPAD4_10754 | -1,231880007 | legumain-like, Expressed effectors |
| WgSM-Oat vs WgSM-Barley | RPAD4_5112 | 1,043289798 | protein 5NUC-like, Expressed effectors |
| WgSM-Oat vs WgSM-Barley | RPAD4_8439 | -1,563317748 | uncharacterized LOC132923934 |
| WgSM-Oat vs WgSM-Barley | RPAD4_13718 | -1,982528303 | glucose dehydrogenase [FAD, quinone]-like |
| WgSM-Oat vs WgSM-Barley | RPAD4_1264 | -1,165779261 | dual specificity protein phosphatase 22-B, Up effectors, Expressed effectors |
| WgSM-Oat vs WgSM-Barley | RPAD4_5454 | -1,818215523 | uncharacterized LOC132918937 |
| WgSM-Oat vs WgSM-Barley | RPAD4_29 | 1,233575861 | major facilitator superfamily domain-containing protein 6-A-like |
| WgSM-Oat vs WgSM-Barley | RPAD4_6357 | -1,093205074 | UDP-glucosyltransferase 2-like |
| WgSM-Oat vs WgSM-Barley | RPAD4_14192 | 1,850945144 | maltase A3-like |
| WgSM-Oat vs WgSM-Barley | RPAD4_10808 | -1,12563749 | uncharacterized LOC132925268 |
| WgSM-Oat vs WgSM-Barley | RPAD4_7926 | -2,363046219 | uncharacterized LOC132919099 |
| WgSM-Oat vs WgSM-Barley | RPAD4_11896 | -1,691054611 | uncharacterized LOC132925750 |
| WgSM-Oat vs WgSM-Barley | RPAD4_12679 | -2,525249525 | heat shock protein 70 A1-like |
| WgSM-Oat vs WgSM-Barley | RPAD4_10495 | -1,611091572 | syntaxin-16-like |
| WgSM-Oat vs WgSM-Barley | RPAD4_13400 | -2,174936567 | heat shock protein 70 A1-like |
| WgSM-Oat vs WgSM-Barley | RPAD4_7135 | -1,118326112 | dynamin-1-like protein |
| WgSM-Oat vs WgSM-Barley | RPAD4_9605 | 1,005380764 | uncharacterized LOC132927366 |
| WgSM-Oat vs WgSM-Barley | RPAD4_12948 | -1,058873173 | neuronal acetylcholine receptor subunit beta-2-like |
| WgSM-Oat vs WgSM-Barley | RPAD4_6731 | -2,072618564 | histone H3, Expressed effectors |
| WgSM-Oat vs WgSM-Barley | RPAD4_14935 | -1,690537384 | uncharacterized LOC132930032 |

| Contrast | Gene_id | logFC | Function |
|---|---|---|---|
| WgSM-Oat vs WgSM-Ryegrass | RPAD4_7192 | 1,695647105 | uncharacterized LOC132921431 |
| WgSM-Oat vs WgSM-Ryegrass | RPAD4_4764 | 3,34531451 | histone H1A, sperm-like |
| WgSM-Oat vs WgSM-Ryegrass | RPAD4_4805 | 1,252296027 | nose resistant to fluoxetine protein 6-like |
| WgSM-Oat vs WgSM-Ryegrass | RPAD4_6869 | 1,142701278 | uncharacterized LOC132922618 |
| WgSM-Oat vs WgSM-Ryegrass | RPAD4_7955 | 2,379780936 | glucose dehydrogenase [FAD, quinone]-like, Expressed effectors |
| WgSM-Oat vs WgSM-Ryegrass | RPAD4_11986 | 1,243841425 | uncharacterized LOC132926493 |
| WgSM-Oat vs WgSM-Ryegrass | RPAD4_12771 | 1,02655756 | uncharacterized LOC132930564 |
| WgSM-Oat vs WgSM-Ryegrass | RPAD4_14316 | 1,08809576 | probable multidrug resistance-associated protein lethal(2)03659 |
| WgSM-Oat vs WgSM-Wheat | RPAD4_12628 | -1,469177018 | uncharacterized LOC132929792 |
| WgSM-Oat vs WgSM-Wheat | RPAD4_5844 | -1,018464499 | probable cytochrome P450 6a13 |
| WgSM-Oat vs WgSM-Wheat | RPAD4_7345 | -1,019391821 | UDP-glucosyltransferase 2-like |
| WgSM-Oat vs WgSM-Wheat | RPAD4_14197 | -1,549586002 | maltase 2-like |
| WgSM-Oat vs WgSM-Wheat | RPAD4_13072 | -1,242880737 | UDP-glucosyltransferase 2-like |
| WgSM-Oat vs WgSM-Wheat | RPAD4_10754 | -1,426212017 | legumain-like, Expressed effectors |
| WgSM-Oat vs WgSM-Wheat | RPAD4_15129 | -1,534317925 | uncharacterized LOC132929383 |
| WgSM-Oat vs WgSM-Wheat | RPAD4_14465 | -1,307975458 | uncharacterized LOC132930328 |
| WgSM-Oat vs WgSM-Wheat | RPAD4_8687 | 4,479489544 | cathepsin B-like, Expressed effectors |
| WgSM-Oat vs WgSM-Wheat | RPAD4_10561 | -1,038242656 | uncharacterized LOC132927497 |
| WgSM-Oat vs WgSM-Wheat | RPAD4_8688 | 3,64239389 | cathepsin B-like, Expressed effectors |
| WgSM-Oat vs WgSM-Wheat | RPAD4_12605 | -1,500355701 | mucin-2-like |
| WgSM-Oat vs WgSM-Wheat | RPAD4_14464 | -1,227343163 | uncharacterized LOC132930276 |
| WgSM-Oat vs WgSM-Wheat | RPAD4_5843 | -1,866588064 | cytochrome P450 6k1-like |
| WgSM-Oat vs WgSM-Wheat | RPAD4_12817 | -1,142295223 | beta-1,3-galactosyltransferase 6-like |
| WgSM-Oat vs WgSM-Wheat | RPAD4_14092 | -1,514252187 | uncharacterized LOC132930687 |
| WgSM-Oat vs WgSM-Wheat | RPAD4_749 | -1,212040064 | cathepsin B-like cysteine proteinase 4, Expressed effectors |
| WgSM-Oat vs WgSM-Wheat | RPAD4_13010 | -2,146208715 | probable cytochrome P450 6a13 |
| WgSM-Oat vs WgSM-Wheat | RPAD4_1327 | -1,373105764 | uncharacterized LOC132917367 |
| WgSM-Oat vs WgSM-Wheat | RPAD4_13318 | -1,035098889 | cuticlin-4-like |
| WgSM-Oat vs WgSM-Wheat | RPAD4_1312 | -1,110021268 | Karl - lipocalin/cytosolic fatty acid-binding protein Karl, Expressed effectors |
| WgSM-Oat vs WgSM-Wheat | RPAD4_10858 | -1,265440822 | uncharacterized LOC132924179 |
| WgSM-Oat vs WgSM-Wheat | RPAD4_10013 | -1,2318049 | uncharacterized LOC132927138 |
| WgSM-Oat vs WgSM-Wheat | RPAD4_7947 | -1,717101043 | myogenesis-regulating glycosidase-like, Expressed effectors |
| WgSM-Oat vs WgSM-Wheat | RPAD4_12750 | -1,045527254 | trehalase-like |
| WgSM-Oat vs WgSM-Wheat | RPAD4_3414 | -1,271780969 | UDP-glucosyltransferase 2-like |
| WgSM-Oat vs WgSM-Wheat | RPAD4_791 | -1,593033311 | major facilitator superfamily domain-containing protein 6-A-like |
| WgSM-Oat vs WgSM-Wheat | RPAD4_13009 | -1,187971308 | probable cytochrome P450 6a13 |
| WgSM-Oat vs WgSM-Wheat | RPAD4_7609 | -1,029739423 | uncharacterized LOC132921815 |
| WgSM-Oat vs WgSM-Wheat | RPAD4_11051 | 1,035017211 | dynein beta chain, ciliary-like |
| WgSM-Oat vs WgSM-Wheat | RPAD4_10557 | -1,289817235 | phenoloxidase 1-like |
| WgSM-Oat vs WgSM-Wheat | RPAD4_3799 | 1,091831228 | uncharacterized LOC132917972 |
| WgSM-Oat vs WgSM-Wheat | RPAD4_6129 | -1,894738009 | UDP-glucosyltransferase 2-like |
| WgSM-Oat vs WgSM-Wheat | RPAD4_10014 | -1,026867061 | uncharacterized LOC132927283 |

| Contrast | Gene_id | logFC | Function |
|---|---|---|---|
| WgSM-Oat vs WgSM-Wheat | RPAD4_750 | -1,002648484 | cathepsin B-like, Expressed effectors |
| WgSM-Oat vs WgSM-Wheat | RPAD4_7198 | 1,07313943 | zinc finger protein 771-like |
| WgSM-Oat vs WgSM-Wheat | RPAD4_11353 | 1,021137943 | uncharacterized LOC132924349 |
| WgSM-Oat vs WgSM-Wheat | RPAD4_11631 | 1,133334863 | NF-kappa-B-activating protein-like |
| WgSM-Oat vs WgSM-Wheat | RPAD4_981 | 1,373366159 | snky - ubiquitin protein ligase sneaky |
| WgSM-Oat vs WgSM-Wheat | RPAD4_7926 | -1,788860053 | uncharacterized LOC132919099 |
| WgSM-Oat vs WgSM-Wheat | RPAD4_9713 | -1,5798014 | uncharacterized LOC132924410 |
| WgSM-Oat vs WgSM-Wheat | RPAD4_12213 | 1,086640055 | zinc finger MYM-type protein 1-like |
| WgSM-Oat vs WgSM-Wheat | RPAD4_4276 | 1,01509952 | uncharacterized LOC132921476 |
| WgSM-Oat vs WgSM-Wheat | RPAD4_10132 | 1,016290241 | cytochrome P450 4C1-like |
| WgSM-Wheat vs WgSM-Barley | RPAD4_14649 | 1,970827432 | germ cell nuclear acidic protein-like |
| WgSM-Wheat vs WgSM-Barley | RPAD4_8218 | 1,26806245 | uncharacterized LOC132922113 |
| WgSM-Wheat vs WgSM-Barley | RPAD4_12628 | 1,677047333 | uncharacterized LOC132929792 |
| WgSM-Wheat vs WgSM-Barley | RPAD4_15129 | 2,048342454 | uncharacterized LOC132929383 |
| WgSM-Wheat vs WgSM-Barley | RPAD4_8837 | 1,372306658 | Osi22 - Osiris 22 |
| WgSM-Wheat vs WgSM-Barley | RPAD4_1980 | -1,016923918 | dentin sialophosphoprotein-like |
| WgSM-Wheat vs WgSM-Barley | RPAD4_569 | 1,869294003 | Not found |
| WgSM-Wheat vs WgSM-Barley | RPAD4_5112 | 1,452881074 | protein 5NUC-like, Expressed effectors |
| WgSM-Wheat vs WgSM-Barley | RPAD4_6887 | 1,652582842 | Osi8 - Osiris 8 |
| WgSM-Wheat vs WgSM-Barley | RPAD4_14465 | 1,475644848 | uncharacterized LOC132930328 |
| WgSM-Wheat vs WgSM-Barley | RPAD4_11631 | -2,168153354 | NF-kappa-B-activating protein-like |
| WgSM-Wheat vs WgSM-Barley | RPAD4_1311 | 2,497726264 | dy - transmembrane protein dusky |
| WgSM-Wheat vs WgSM-Barley | RPAD4_13395 | 1,03603238 | peroxidase-like, Up effectors, Salivary proteins, Expressed effectors |
| WgSM-Wheat vs WgSM-Barley | RPAD4_7609 | 1,455290422 | uncharacterized LOC132921815 |
| WgSM-Wheat vs WgSM-Barley | RPAD4_7610 | 1,281672802 | uncharacterized LOC132922084 |
| WgSM-Wheat vs WgSM-Barley | RPAD4_12689 | 1,426596003 | ichor-like |
| WgSM-Wheat vs WgSM-Barley | RPAD4_13318 | 1,221438895 | cuticlin-4-like |
| WgSM-Wheat vs WgSM-Barley | RPAD4_7198 | -1,844297748 | zinc finger protein 771-like |
| WgSM-Wheat vs WgSM-Barley | RPAD4_5776 | 1,335168382 | uncharacterized LOC132920535 |
| WgSM-Wheat vs WgSM-Barley | RPAD4_10559 | 1,011798934 | phenoloxidase 1-like |
| WgSM-Wheat vs WgSM-Barley | RPAD4_12844 | 1,251628095 | agrin-like, Expressed effectors |
| WgSM-Wheat vs WgSM-Barley | RPAD4_10620 | -1,062346177 | uncharacterized LOC132926451 |
| WgSM-Wheat vs WgSM-Barley | RPAD4_12605 | 1,489326772 | mucin-2-like |
| WgSM-Wheat vs WgSM-Barley | RPAD4_11955 | 7,059766212 | uncharacterized LOC132925785 |
| WgSM-Wheat vs WgSM-Barley | RPAD4_1327 | 1,481764238 | uncharacterized LOC132917367 |
| WgSM-Wheat vs WgSM-Barley | RPAD4_29 | 1,488835343 | major facilitator superfamily domain-containing protein 6-A-like |
| WgSM-Wheat vs WgSM-Barley | RPAD4_6005 | -1,63648087 | receptor expression-enhancing protein 5-like |
| WgSM-Wheat vs WgSM-Barley | RPAD4_10014 | 1,463109544 | uncharacterized LOC132927283 |
| WgSM-Wheat vs WgSM-Barley | RPAD4_8217 | 1,910216303 | uncharacterized LOC132922903 |
| WgSM-Wheat vs WgSM-Barley | RPAD4_2648 | 1,587248391 | Cad89D - cadherin 89D |
| WgSM-Wheat vs WgSM-Barley | RPAD4_6437 | -1,058814328 | lisH domain-containing protein ARMC9-like |
| WgSM-Wheat vs WgSM-Barley | RPAD4_7613 | 1,035946101 | uncharacterized LOC132921612 |
| WgSM-Wheat vs WgSM-Barley | RPAD4_9198 | 2,279273096 | uncharacterized LOC132924308 |

| Contrast | Gene_id | logFC | Function |
|---|---|---|---|
| WgSM-Wheat vs WgSM-Barley | RPAD4_9522 | -1,71862192 | dynein axonemal heavy chain 3-like |
| WgSM-Wheat vs WgSM-Barley | RPAD4_12626 | 1,355207656 | uncharacterized LOC132929682 |
| WgSM-Wheat vs WgSM-Barley | RPAD4_9065 | -1,054282658 | uncharacterized LOC132924433 |
| WgSM-Wheat vs WgSM-Barley | RPAD4_4272 | -1,112352641 | noggin-1-like, Expressed effectors |
| WgSM-Wheat vs WgSM-Barley | RPAD4_10808 | -1,346790065 | uncharacterized LOC132925268 |
| WgSM-Wheat vs WgSM-Barley | RPAD4_3882 | -1,138728069 | uncharacterized LOC132927309 |
| WgSM-Wheat vs WgSM-Barley | RPAD4_4432 | 1,297942896 | uncharacterized LOC132921531 |
| WgSM-Wheat vs WgSM-Barley | RPAD4_6265 | -8,116331463 | histone H3, Expressed effectors |
| WgSM-Wheat vs WgSM-Barley | RPAD4_10013 | 1,256114558 | uncharacterized LOC132927138 |
| WgSM-Wheat vs WgSM-Barley | RPAD4_10824 | -1,139957093 | uncharacterized LOC132927739 |
| WgSM-Wheat vs WgSM-Barley | RPAD4_8990 | -1,257721981 | major antigen-like |
| WgSM-Wheat vs WgSM-Barley | RPAD4_7906 | 1,127285281 | cell-death-related nuclease 7-like, Expressed effectors |
| WgSM-Wheat vs WgSM-Barley | RPAD4_12817 | 1,095984946 | beta-1,3-galactosyltransferase 6-like |
| WgSM-Wheat vs WgSM-Barley | RPAD4_610 | -4,13129053 | cuticle protein 16.5-like, Expressed effectors |
| WgSM-Wheat vs WgSM-Barley | RPAD4_14621 | 1,054391775 | alpha-1,3-mannosyl-glycoprotein 4-beta-N-acetylglucosaminyltransferase B-like |
| WgSM-Wheat vs WgSM-Barley | RPAD4_1056 | 1,243228479 | carboxypeptidase B-like, Expressed effectors |
| WgSM-Wheat vs WgSM-Barley | RPAD4_5245 | 1,122385485 | carbonic anhydrase 2-like, Up effectors, Expressed effectors |
| WgSM-Wheat vs WgSM-Barley | RPAD4_14464 | 1,081524832 | uncharacterized LOC132930276 |
| WgSM-Wheat vs WgSM-Barley | RPAD4_12602 | 1,620156005 | uncharacterized LOC132928706 |
| WgSM-Wheat vs WgSM-Barley | RPAD4_3837 | 1,427503841 | uncharacterized LOC132917960 |
| WgSM-Wheat vs WgSM-Barley | RPAD4_7703 | -1,10876984 | Kif3C - Kinesin family member 3C |
| WgSM-Wheat vs WgSM-Barley | RPAD4_13118 | -1,136365844 | luciferin 4-monooxygenase-like |
| WgSM-Wheat vs WgSM-Barley | RPAD4_3476 | 1,512687656 | polyprenol reductase |
| WgSM-Wheat vs WgSM-Barley | RPAD4_9731 | -1,371250612 | cilia- and flagella-associated protein 52-like |
| WgSM-Wheat vs WgSM-Barley | RPAD4_12194 | -1,082021555 | uncharacterized LOC132925991 |
| WgSM-Wheat vs WgSM-Barley | RPAD4_2931 | 1,786082002 | phospholipase A1-like |
| WgSM-Wheat vs WgSM-Barley | RPAD4_6995 | -1,385335885 | uncharacterized LOC132921433 |
| WgSM-Wheat vs WgSM-Barley | RPAD4_10074 | 1,181603027 | uncharacterized LOC132927448 |
| WgSM-Wheat vs WgSM-Barley | RPAD4_10673 | -1,065198541 | uncharacterized LOC132926772 |
| WgSM-Wheat vs WgSM-Barley | RPAD4_10932 | -1,126220053 | short transient receptor potential channel 4-like |
| WgSM-Wheat vs WgSM-Barley | RPAD4_12630 | 1,174842335 | uncharacterized LOC132929790 |
| WgSM-Wheat vs WgSM-Barley | RPAD4_14092 | 1,431338795 | uncharacterized LOC132930687 |
| WgSM-Wheat vs WgSM-Barley | RPAD4_6880 | 2,67362073 | uncharacterized LOC132922347 |
| WgSM-Wheat vs WgSM-Barley | RPAD4_2545 | -1,320856085 | dynein axonemal heavy chain 7-like |
| WgSM-Wheat vs WgSM-Barley | RPAD4_6864 | 1,284306055 | Osi24 - Protein Osi24, Expressed effectors |
| WgSM-Wheat vs WgSM-Barley | RPAD4_10716 | -1,502120775 | trichohyalin-like |
| WgSM-Wheat vs WgSM-Barley | RPAD4_8439 | -1,470494855 | uncharacterized LOC132923934 |
| WgSM-Wheat vs WgSM-Barley | RPAD4_12693 | -1,18520518 | uncharacterized LOC132930145 |
| WgSM-Wheat vs WgSM-Barley | RPAD4_13247 | 1,80995767 | neuroendocrine convertase 1-like |
| WgSM-Wheat vs WgSM-Barley | RPAD4_3708 | 1,981544817 | uncharacterized LOC132916989 |
| WgSM-Wheat vs WgSM-Barley | RPAD4_13435 | -1,223464982 | uncharacterized LOC132930167 |
| WgSM-Wheat vs WgSM-Barley | RPAD4_1688 | -1,147455605 | nuclear receptor ROR-alpha B-like |

| Contrast | Gene_id | logFC | Function |
|---|---|---|---|
| WgSM-Wheat vs WgSM-Barley | RPAD4_3172 | -1,360463058 | uncharacterized LOC132917842 |
| WgSM-Wheat vs WgSM-Barley | RPAD4_6945 | 1,335517082 | uncharacterized LOC132921646 |
| WgSM-Wheat vs WgSM-Barley | RPAD4_12834 | 1,421142735 | uncharacterized LOC132928690 |
| WgSM-Wheat vs WgSM-Barley | RPAD4_14368 | -1,220084005 | adenylate kinase 9-like |
| WgSM-Wheat vs WgSM-Barley | RPAD4_10867 | -1,111453622 | serine-aspartate repeat-containing protein I-like |
| WgSM-Wheat vs WgSM-Barley | RPAD4_981 | -1,856487962 | snky - ubiquitin protein ligase sneaky |
| WgSM-Wheat vs WgSM-Barley | RPAD4_11128 | -1,302677078 | keratin-associated protein 10-7-like |
| WgSM-Wheat vs WgSM-Barley | RPAD4_11578 | -1,189583945 | RNA polymerase-associated protein CTR9 homolog |
| WgSM-Wheat vs WgSM-Barley | RPAD4_7683 | 1,061268062 | uncharacterized LOC132922088 |
| WgSM-Wheat vs WgSM-Barley | RPAD4_11051 | -1,031335523 | dynein beta chain, ciliary-like |
| WgSM-Wheat vs WgSM-Barley | RPAD4_10846 | -1,100767971 | uncharacterized LOC132925215 |
| WgSM-Wheat vs WgSM-Barley | RPAD4_14935 | -2,397399489 | uncharacterized LOC132930032 |
| WgSM-Wheat vs WgSM-Barley | RPAD4_15044 | -1,272726053 | testis-specific gene A8 protein-like |
| WgSM-Wheat vs WgSM-Barley | RPAD4_2462 | -1,441179167 | Gr63a - Gustatory receptor 63a |
| WgSM-Wheat vs WgSM-Barley | RPAD4_9497 | -1,32254652 | Arl6 - ADP ribosylation factor-like 6 |
| WgSM-Wheat vs WgSM-Barley | RPAD4_352 | 1,079067872 | glutathione S-transferase-like |
| WgSM-Wheat vs WgSM-Barley | RPAD4_5843 | 1,309184005 | cytochrome P450 6k1-like |
| WgSM-Wheat vs WgSM-Barley | RPAD4_13413 | 1,067967959 | eukaryotic translation initiation factor 4E-like |
| WgSM-Wheat vs WgSM-Barley | RPAD4_12650 | -1,133675111 | uncharacterized LOC132928673 |
| WgSM-Wheat vs WgSM-Barley | RPAD4_13130 | -1,00173658 | NFX1-type zinc finger-containing protein 1-like |
| WgSM-Wheat vs WgSM-Barley | RPAD4_379 | -1,029403357 | protein maelstrom 1-like |
| WgSM-Wheat vs WgSM-Barley | RPAD4_12509 | -1,071659092 | uncharacterized LOC132926261 |
| WgSM-Wheat vs WgSM-Barley | RPAD4_754 | -1,1257185 | uncharacterized LOC132931762 |
| WgSM-Wheat vs WgSM-Barley | RPAD4_7945 | -1,133060739 | uncharacterized LOC132923000 |
| WgSM-Wheat vs WgSM-Barley | RPAD4_11761 | -1,225278577 | uncharacterized LOC132925677 |
| WgSM-Wheat vs WgSM-Barley | RPAD4_5292 | -1,327333486 | uncharacterized LOC132922308 |
| WgSM-Wheat vs WgSM-Barley | RPAD4_4913 | -1,155151475 | uncharacterized LOC132922619 |
| WgSM-Wheat vs WgSM-Barley | RPAD4_13025 | -1,01384116 | glycerol kinase-like |
| WgSM-Wheat vs WgSM-Barley | RPAD4_7510 | 1,113125966 | uncharacterized LOC132919704 |
| WgSM-Wheat vs WgSM-Barley | RPAD4_10748 | 1,175128808 | uncharacterized LOC132927446 |
| WgSM-Wheat vs WgSM-Barley | RPAD4_8310 | 1,55209104 | glucose dehydrogenase [FAD, quinone]-like, Expressed effectors |
| WgSM-Wheat vs WgSM-Barley | RPAD4_11896 | -1,355190661 | uncharacterized LOC132925750 |
| WgSM-Wheat vs WgSM-Barley | RPAD4_9077 | -1,15040989 | tubulin beta chain-like |
| WgSM-Wheat vs WgSM-Barley | RPAD4_3897 | -1,1777952 | uncharacterized LOC132918684 |
| WgSM-Wheat vs WgSM-Barley | RPAD4_11419 | -1,266743796 | E3 ubiquitin-protein ligase mind-bomb-like |
| WgSM-Wheat vs WgSM-Barley | RPAD4_14287 | 1,350219552 | glucose dehydrogenase [FAD, quinone]-like, Salivary proteins |
| WgSM-Wheat vs WgSM-Barley | RPAD4_3562 | -1,051660829 | uncharacterized LOC132932115 |
| WgSM-Wheat vs WgSM-Barley | RPAD4_6890 | 1,200035023 | uncharacterized LOC132920415 |
| WgSM-Wheat vs WgSM-Barley | RPAD4_11360 | -2,298116758 | THAP domain-containing protein 2-like, Expressed effectors |

**Supplementary table 4. Comparative metabolite profiles of leaf and stem phloem of Poaceae hosts of *R. padi* (wheat. oat. barley. and ryegrass)**. Data were analyzed using one-way ANOVA with Tukey's *post hoc* test for homogeneous datasets and Kruskal–Wallis with Dunn's *post hoc* test (Bonferroni correction) for non-homogeneous datasets. ND = not detected. Lowercase letters in superscript adjacent to compound concentration mean values represent significant differences ($p \leq 0.05$) determined using *post hoc* tests. Asterisks adjacent to amino acid names indicate essential amino acids.

| **Compound** | | | **Compound concentration (mean ± SE in nmoles mL$^{-1}$ phloem)** | | | | **Statistics** |
|---|---|---|---|---|---|---|---|
| **Class** | **Chemical nature** | **Name** | **Wheat** | **Oat** | **Barley** | **Ryegrass** | |
| Amino acid | Polar basic | Histidine* | 28.4$^{b}$ ± 2.62 | 27.57$^{b}$ ± 3.94 | 18.71$^{a}$ ± 2.98 | 16.72$^{a}$ ± 2.51 | $\chi^2$ = 9.05. *p* = 0.03 |
| | | Arginine* | 76.1$^{c}$ ± 7.98 | 60.32$^{bc}$ ± 8.11 | 33.8$^{a}$ ± 2.96 | 47.22$^{ab}$ ± 5.66 | $\chi^2$ = 17.51. *p* = 0.0006 |
| | | Lysine* | 139.99$^{b}$ ± 14.31 | 151.77$^{b}$ ± 31.05 | 74.75$^{a}$ ± 8.35 | 57.64$^{a}$ ± 5.31 | $\chi^2$ = 21.07. *p* = 1.00E-04 |
| | Polar amide containing | Asparagine | 46.84$^{ab}$ ± 4.63 | 94.63$^{c}$ ± 15.47 | 30.91$^{a}$ ± 3.43 | 83.54$^{bc}$ ± 15.57 | $\chi^2$ = 19.09. *p* = 0.0003 |
| | | Glutamine | 159.12$^{b}$ ± 14.5 | 263.3$^{bc}$ ± 35.7 | 105.36$^{a}$ ± 10.81 | 543.36$^{c}$ ± 88.08 | $\chi^2$ = 28.23. *p* = 3.25E-06 |
| | Polar acidic | Aspartate | 532.04$^{b}$ ± 54.4 | 545.96$^{b}$ ± 83.24 | 317.75$^{a}$ ± 34.85 | 693.49$^{b}$ ± 61.73 | $\chi^2$ = 14.67. *p* = 0.002 |
| | | Glutamate | 126.26$^{a}$ ± 23.39 | 312.63$^{b}$ ± 46.26 | 130.68$^{a}$ ± 14.72 | 640.96$^{b}$ ± 77.53 | $\chi^2$ = 28.17. *p* = 3.35E-06 |
| | Polar hydroxyl containing | Serine | 322.06$^{b}$ ± 32.75 | 316.68$^{ab}$ ± 51.72 | 197.25$^{a}$ ± 21.67 | 593.12$^{c}$ ± 66.19 | $\chi^2$ = 19.62. *p* = 0.0002 |
| | | Threonine* | 110.16$^{ab}$ ± 9.64 | 158.99$^{bc}$ ± 25.59 | 85.01$^{a}$ ± 8.81 | 238.51c ± 26.04 | $\chi^2$ = 17.75. *p* = 0.0005 |
| | Polar aromatic hydroxyl containing | Tyrosine | 55.47$^{b}$ ± 5.85 | 81.61$^{b}$ ± 14.3 | 30.07$^{a}$ ± 2.97 | 34.39$^{a}$ ± 3.39 | $\chi^2$ = 21.48. *p* = 8.37E-05 |
| | Nonpolar aromatic | Phenylalanine* | 71.9$^{bc}$ ± 6.6 | 92.71$^{c}$ ± 18.38 | 32.5$^{a}$ ± 3.1 | 50.62$^{ab}$ ± 6.21 | $\chi^2$ = 19.15. *p* = 3.00E-04 |
| | | Tryptophane* | 16.34$^{b}$ ± 1.65 | 25.1$^{b}$ ± 4.63 | 16.47$^{b}$ ± 1.65 | 4.8$^{a}$ ± 1.15 | $\chi^2$ = 23.2. *p* = 3.66E-05 |
| | Nonpolar aliphatic | Glycine | 46.8$^{b}$ ± 5.06 | 56.94$^{b}$ ± 11.05 | 28.79$^{a}$ ± 3.21 | 235.4$^{ab}$ ± 192.68 | $\chi^2$ = 8.56. *p* = 0.04 |
| | | Alpha-Alanine | 336$^{a}$ ± 30.89 | 673.66$^{b}$ ± 105.61 | 288.93$^{a}$ ± 28.88 | 561.46$^{b}$ ± 45.33 | $\chi^2$ = 21.65. *p* = 7.70E-05 |
| | | Proline | 23.78$^{a}$ ± 2.6 | 47.28$^{b}$ ± 8.09 | 30.71$^{a}$ ± 2.36 | 47.6$^{b}$ ± 4.78 | $\chi^2$ = 14.77. *p* = 0.002 |
| | | Methionine* | 33.96$^{b}$ ± 3.86 | 40.77$^{b}$ ± 7.02 | 13.04$^{a}$ ± 1.18 | 17.07$^{a}$ ± 3.08 | $\chi^2$ = 23.57. *p* = 3.07E-05 |
| | | Valine* | 104.44$^{b}$ ± 8.87 | 119.48$^{ab}$ ± 19.44 | 75.36$^{a}$ ± 6.98 | 102.35$^{ab}$ ± 9.64 | $\chi^2$ = 5.55. *p* = 1.36E-01 |
| | | Isoleucine* | 45.29 ± 4.15 | 43.59 ± 7.23 | 35.11 ± 3.2 | 32.54 ± 2.95 | $\chi^2$ = 4.3. *p* = 0.2 |
| | | Leucine* | 93.09$^{b}$ ± 9.33 | 117.69$^{b}$ ± 24.71 | 44.53$^{a}$ ± 4.45 | 53.38$^{a}$ ± 4.9 | $\chi^2$ = 19.68. *p* = 2.00E-04 |
| Sugar and sugar alcohol | Monosaccharide | Glucose | 677.41$^{a}$ ± 83.77 | 1597.76$^{b}$ ± 103.26 | 1635.94$^{b}$ ± 118.83 | 544.7$^{a}$ ± 46.62 | $\chi^2$ = 28.45. *p* = 2.9E-06 |
| | | Fructose | 621.8$^{a}$ ± 94.19 | 1551.35$^{b}$ ± 121.87 | 1294.7$^{b}$ ± 132.93 | 708.75$^{a}$ ± 75.96 | $\chi^2$ = *23.57*. *p* = 3.07E-05 |
| | | Galactose | 48.41$^{a}$ ± 2.75 | 49.37$^{a}$ ± 2.58 | 74.86$^{b}$ ± 5.44 | 56.27$^{a}$ ± 5.5 | $\chi^2$ = 12.85. *p* = 0.005 |
| | | Arabinose | 15.41$^{ab}$ ± 0.61 | 14.82$^{ab}$ ± 0.33 | 13.59$^{a}$ ± 0.33 | 16.14$^{b}$ ± 0.64 | *df* = 3. *F* = 4.25. p = 0.01 |
| | | Xylose | 15.11$^{a}$ ± 0.22 | 16$^{ab}$ ± 0.25 | 16.51$^{b}$ ± 0.26 | 16.92$^{b}$ ± 0.35 | *df* = 3. *F* = 7.44. p = 0.0005 |
| | Disaccharide | Sucrose | 21.29$^{a}$ ± 0.28 | 620.95$^{b}$ ± 85.29 | 22.11$^{a}$ ± 0.45 | 24.32$^{a}$ ± 2.89 | $\chi^2$ = 23.25. *p* = 3.80E-05 |

| Compound | | | Compound concentration (mean ± SE in nmoles $mL^{-1}$ phloem) | | | | Statistics |
|---|---|---|---|---|---|---|---|
| Class | Chemical nature | Name | Wheat | Oat | Barley | Ryegrass | |
| | | Maltose | 47.57$^{a}$ ± 0.26 | 51.52$^{b}$ ± 0.66 | 49.24$^{b}$ ± 0.55 | 47.54$^{a}$ ± 0.18 | $\chi^2$ = 23.83. *p* = 2.70E-05 |
| | | Melibiose | 34.58$^{a}$ ± 0.14 | 34.47$^{a}$ ± 0.11 | 39.95$^{b}$ ± 0.6 | 37.89$^{b}$ ± 0.92 | $\chi^2$ = 29.59. *p* = 1.67E-06 |
| | Sugar alcohol | Mannitol | 2.91 ± 1.15 | 7.53 ± 1.62 | ND | 6.35 ± 1.55 | |
| | | Dulcitol | 0.82 ± 0.22 | 0.64 ± 0.31 | ND | ND | |
| | | Myoinositol | 20.56$^{a}$ ± 4.01 | 81.88$^{c}$ ± 5.13 | 42.09$^{b}$ ± 3.04 | 52.42$^{b}$ ± 3.2 | *df* = 3. *F* = 37.79. p = 3.24E-11 |
| | | Galactinol | 14.34$^{a}$ ± 0.23 | 13.67$^{a}$ ± 0.11 | 16.45$^{b}$ ± 0.7 | 13.74$^{a}$ ± 0.23 | $\chi^2$ = 19.56. *p* = 2.00E-04 |
| Organic acid | Hydroxy carboxylic acid | Glycolic acid | 7.74$^{a}$ ± 0.24 | 8.35$^{a}$ ± 0.3 | 7.31$^{a}$ ± 0.33 | 9.71$^{b}$ ± 0.31 | *df* = 3. *F* = 11.17. p = 2.50E-5 |
| | Hydroxy acid | Glyceric acid | 30.74 ± 2.21 | 33.49 ± 2.19 | 27.18 ± 2.26 | 31.15 ± 2.85 | *df* = 3. *F* = 1.062. p = 0.38 |
| | Carboxylic acid | Succinic acid | 26.05$^{a}$ ± 2.64 | 106.31$^{b}$ ± 7.94 | 22.64$^{a}$ ± 1.94 | 66.5$^{b}$ ± 7.96 | $\chi^2$ = 30.32. *p* = 1.18E-06 |
| | | Fumaric acid | 10.29$^{a}$ ± 0.24 | 15.3$^{b}$ ± 0.43 | 11.33$^{a}$ ± 0.32 | 18.08$^{b}$ ± 1.45 | $\chi^2$ = 29.3. *p* = 1.93E-06 |
| | | Malic acid | 127.16$^{a}$ ± 22.5 | 307.52$^{b}$ ± 22.28 | 359.82$^{b}$ ± 42.11 | 589.39$^{b}$ ± 102.65 | $\chi^2$ = 31.73. *p* = 7.40E-05 |
| | | Citric acid | 54.81$^{a}$ ± 5.63 | 98.3$^{ab}$ ± 7.93 | 62.59$^{a}$ ± 3.64 | 132.15$^{b}$ ± 20.05 | *df* = 3. *F* = 8.94. p = 0.0001 |
| | Cyclic polyhydroxy carboxylic acid | Quinic acid | 31.68$^{a}$ ± 3.44 | 97.44$^{b}$ ± 7.93 | 128.54$^{b}$ ± 12.57 | 464.14$^{c}$ ± 44.5 | $\chi^2$ = 33.99. *p* = 1.90E-07 |

**Supplementary table 5. Results of Cox proportional hazards models testing the effects of acidic water and quinic acid supplementation on survival of WlSM.** The table reports regression coefficients (coef), hazard ratios (exp(coef)), standard errors (se(coef)), z statistics (z), and associated *P*-values (Pr(>\|z\|)) for each pairwise contrast. Hazard ratios greater than 1 indicate an increased risk of death relative to the reference treatment, whereas values below 1 indicate a reduced risk.

| Morph | Contrast | coef | exp(coef) | se(coef) | z | Pr(>\|z\|) |
|---|---|---|---|---|---|---|
| WlSM | Quinic acid vs acidic water | -20.298 | 1.530e-09 | 9971.793 | -0.002 | 0.998 |

**Supplementary table 6. Post hoc comparisons of cumulative fecundity among aphid morphs under acidic water and quinic acid treatments.** The table presents pairwise contrasts of estimated marginal means derived from linear mixed-effects models testing the effects of morph, treatment (acidic water and quinic acid), and their interaction on cumulative fecundity. For each contrast, the estimated difference (estimate), standard error (SE), degrees of freedom (df), *t* statistic (*t*.ratio), and associated *P*-value (*p*.value) are reported. *P*-values were adjusted for multiple testing using Tukey's method. Positive estimates indicate higher fecundity in the first level of the contrast relative to the second.

| Morph | contrast | estimate | SE | df | t.ratio | p.value |
|---|---|---|---|---|---|---|
| WlSM | Quinic acid vs acidic water | -10,385 | 0,703 | 217,005 | -14,765 | 1,287e-34 |
